# Non-Invasive Plasma Glycomic and Metabolic Biomarkers of Post-treatment Control of HIV

**DOI:** 10.1101/2020.11.11.378174

**Authors:** Leila B. Giron, Clovis S. Palmer, Qin Liu, Xiangfan Yin, Emmanouil Papasavvas, Mohammad Damra, Aaron R. Goldman, Hsin-Yao Tang, Rowena Johnston, Karam Mounzer, Jay R. Kostman, Pablo Tebas, Alan Landay, Luis J. Montaner, Jeffrey M. Jacobson, Jonathan Z. Li, Mohamed Abdel-Mohsen

**Affiliations:** The Wistar Institute, Philadelphia, PA, 19104, USA; The Burnet Institute, Melbourne, Victoria, 3004, Australia; Department of Infectious Diseases, Monash University, Melbourne, Victoria, 3004, Australia; amfAR, The Foundation for AIDS Research, New York, New York, USA; Philadelphia FIGHT, Philadelphia, PA, 19107, USA; University of Pennsylvania, Philadelphia, PA, 19104, USA; Rush University, Chicago, IL, 60612, USA; Case Western Reserve University School of Medicine, Cleveland, Ohio, 44106, USA; Department of Medicine, Brigham and Women’s Hospital, Harvard Medical School, Boston, MA 02115, USA

**Keywords:** HIV persistence, HIV Cure, HIV Rebound, Analytic Treatment Interruption, Post-treatment control, Glycomic, Metabolomic, L-Glutamic acid, pyruvic acid, Galactose, Fucose

## Abstract

Non-invasive biomarkers that predict HIV remission after antiretroviral therapy (ART) interruption are urgently needed. Such biomarkers can improve the safety of analytic treatment interruption (ATI) and provide mechanistic insights into the pathways involved in post-ART HIV control. We identified plasma glycomic and metabolic signatures of time-to-viral-rebound and probability-of-viral-rebound using samples from two independent cohorts. These samples include a large number of post-treatment controllers, a rare population demonstrating sustained virologic suppression after ART-cessation. The signatures remained significant after adjusting for key demographic and clinical confounders. We also confirmed a mechanistic link between biomarkers and HIV latency reactivation and myeloid inflammation *in vitro*. Finally, machine learning algorithms selected sets of biomarkers that predict time-to-viral-rebound with 74-76% capacity and probability-of-viral-rebound with 97.5% capacity. In summary, we fill a major gap in HIV cure research by identifying non-invasive biomarkers, with potential functional significance, that predict duration and probability of viral remission after treatment interruption.

## INTRODUCTION

Several therapeutic strategies are being tested in clinical trials to reduce the size of HIV reservoirs to a point where virologic control can be achieved without antiretroviral therapy (ART).^1^ The success of these strategies depends on the capacity to determine if potential interventions have made a meaningful impact on the HIV reservoir, i.e. if they have extended the likely period of ART-free remission following treatment discontinuation. Because current technologies are unable to measure the impact of interventions on the total body burden of HIV, HIV cure-focused clinical trials rely on the inclusion of an analytic treatment interruption (ATI) as the only definitive approach to evaluate the effectiveness of interventions.^2-4^ However, this approach is costly, cumbersome, and poses some risk to both study participants and the community. These realities highlight the urgent need for biomarkers that can accurately predict time-to-viral-rebound after treatment interruption and can be leveraged to guide clinical decision making. Such predictive biomarkers could be used to improve the safety of ATIs and accelerate the development of an HIV cure by providing a means for selecting only the most promising therapies for testing by ATIs.^5^ These biomarkers could also provide mechanistic insights into the molecular and biochemical pathways involved in post-ART control of HIV.

In the last few years, a small number of immunophenotypic and virologic measurements have been associated with time-to-viral-rebound. Levels of exhaustion markers on CD4^+^ T cells, measured pre-ART, correlated with time-to-rebound.^6^ However, these measures fail as biomarkers when assessed during ART.^6^ Levels of cell-associated HIV DNA^7^ and RNA,^8,9^ as well as features of plasmacytoid dendritic cells,^10^ during ART, correlate with viral rebound after ART cessation, however, the correlations are generally modest. Thus, as of now, there are no sufficiently reliable or validated biomarkers that can be leveraged to guide clinical decision making.

While the majority of HIV-infected individuals experience rapid viral rebound after ART interruption,^8^ a rare population of individuals, termed post-treatment controllers (PTCs), demonstrate sustained virologic suppression for several months to years after ART cessation.^11-13^ The mechanisms underlying viral control in these individuals are not completely understood. Nonetheless, they represent a clinically relevant model for viral control post-ART.^14,15^ The existence of these individuals with this phenotype raises the question: is it possible to define a set of biomarkers that can predict the probability-of-viral-rebound after potentially successful intervention (i.e., the likelihood to achieve a PTC phenotype after ART cessation)? These biomarkers can also provide critical insights into the mechanisms that underlie this clinically-relevant and desirable phenotype.

We have been taking advantage of work in the emerging fields of glycomics and metabolomics to identify highly robust, host-specific plasma biomarkers that can predict the duration and probability of viral remission after treatment interruption. Plasma glycoproteins (including antibodies; immunoglobulin G (IgGs)) and plasma metabolites enter the circulation from tissues through active secretion or leakage. Therefore, their levels and chemical characteristics can reflect the overall status of multiple organs, making them excellent candidates for biomarker discovery. Indeed, glycomic features in total plasma and on IgG have been identified as biomarkers for inflammatory bowel disease, cancer, and diabetes.^16-21^ In addition, glycans on circulating glycoproteins have functional significance, as they play essential roles in mediating immunological functions, including antibody-dependent cell-mediated cytotoxicity (ADCC) and pro- and anti-inflammatory activities.^22-25^ Similarly, plasma metabolites have been investigated as diagnostic and prognostic biomarkers in several diseases such as heart disease,^26^ hepatitis,^27^ Alzheimer’s disease,^28^ and cancer.^29,30^ Similar to plasma glycans, plasma metabolites are biologically active molecules that function to regulate critical immunological responses, including inflammatory responses.^31-34^

In a recent pilot study,^35^ we identified several plasma glycomic structures whose pre-ATI levels associate with delayed viral rebound after ART discontinuation. These were the digalactosylated glycans on bulk IgG, called G2, as well as fucose (total and core) and *N*-Acetylglucosamine (GlcNac) on total plasma glycoproteins.^35^ However, that study was a small pilot and did not explicitly address the potentially confounding effects of age, gender, ethnicity, duration-on-ART, time of ART initiation (treatment at early vs. chronic stage of infection), or pre-ATI CD4 count.

In this current study, we first extended our biomarker discovery by applying metabolomic analysis on one of the two cohorts used in the pilot.^35^ This was a cohort of 24 HIV-infected, ART-suppressed individuals who had participated in an open-ended ATI study without concurrent immunomodulatory agents. Our metabolomic analysis identified a select set of metabolites whose pre-ATI levels associate with time-to-viral-rebound. These metabolites belong to metabolic pathways known to impact inflammatory responses. We confirmed the direct functional impact of some of these metabolites on latent HIV reactivation and/or macrophage inflammation *in vitro*. We then profiled both the plasma glycome and metabolome of a large cohort of 74 HIV-infected, ART-suppressed individuals who underwent ATI during several AIDS Clinical Trials Group (ACTG) clinical trials. This cohort contains 27 PTCs and 47 post-treatment non-controllers (NCs). Using this cohort, we confirmed the utility of a set of plasma glycans and metabolites to predict time-to-viral-rebound and probability-of-viral-rebound even after adjusting for several potential demographic and clinical confounders. Finally, using machine learning models, we combined this set of biomarkers into two multivariate models: a model that predicts time-to-viral-rebound with 74-76% capacity; and a model that predicts probability-of-viral-rebound (PVR score) with 97.5% capacity. Together, we fill a major gap in HIV cure research by identifying plasma non-invasive biomarkers, with potential functional significance, that predict duration and probability of viral remission after treatment interruption.

## RESULTS

### Characteristics of study cohorts

In this study, we employed two ATI cohorts: 1) The Philadelphia cohort: a cohort of 24 HIV-infected individuals on suppressive ART who underwent an open-ended ATI.^35,36^ This cohort had a wide distribution of viral rebound times (14 to 119 days; median=28; **Supplementary Table 1)**.^35^ Importantly, this cohort underwent ATI without concurrent immunomodulatory agents that might confound our signatures at the initial discovery phase.^35,36^ 2) The AIDS Clinical Trial Group (ACTG) cohort: a cohort combining 74 participants 80 from six ACTG ATI studies (ACTG 371,^37^ A5024,^38^ A5068,^39^ A5170,^40^ A5187,^41^ and A5197^42^), tested or not the efficacy of different HIV vaccines and interleukin-2 (IL-2) treatment. These six ATI studies included 567 participants and identified 27 PTCs out of these participants. Our ACTG cohort included all 27 PTCs and 47 matched NCs from the same studies. The definition of post-treatment control was: remaining off ART for ≥24 weeks post-ATI with VL ≤400 copies for at least 2/3 of time points; had no ART in the plasma; and had no evidence of spontaneous control pre-ART. The remaining 47 were non-controllers (NCs) who rebounded before meeting PTC criteria.^43-45^ The PTC and NC groups within the ACTG cohort are matched for gender, age, ethnicity, % treated during early infection, ART duration and pre-ATI CD4 count (**Table 1** and **Supplementary Figure 1)**. Notably, the combined studies within the ACTG cohort reflect six ATI clinical trials where individuals received or not different HIV vaccines and/or immunotherapies.^37-42^ This important feature of this cohort allows for identifying/validating markers that predict duration and probability of viral remission independent of potential interventions.

**Table 1.**
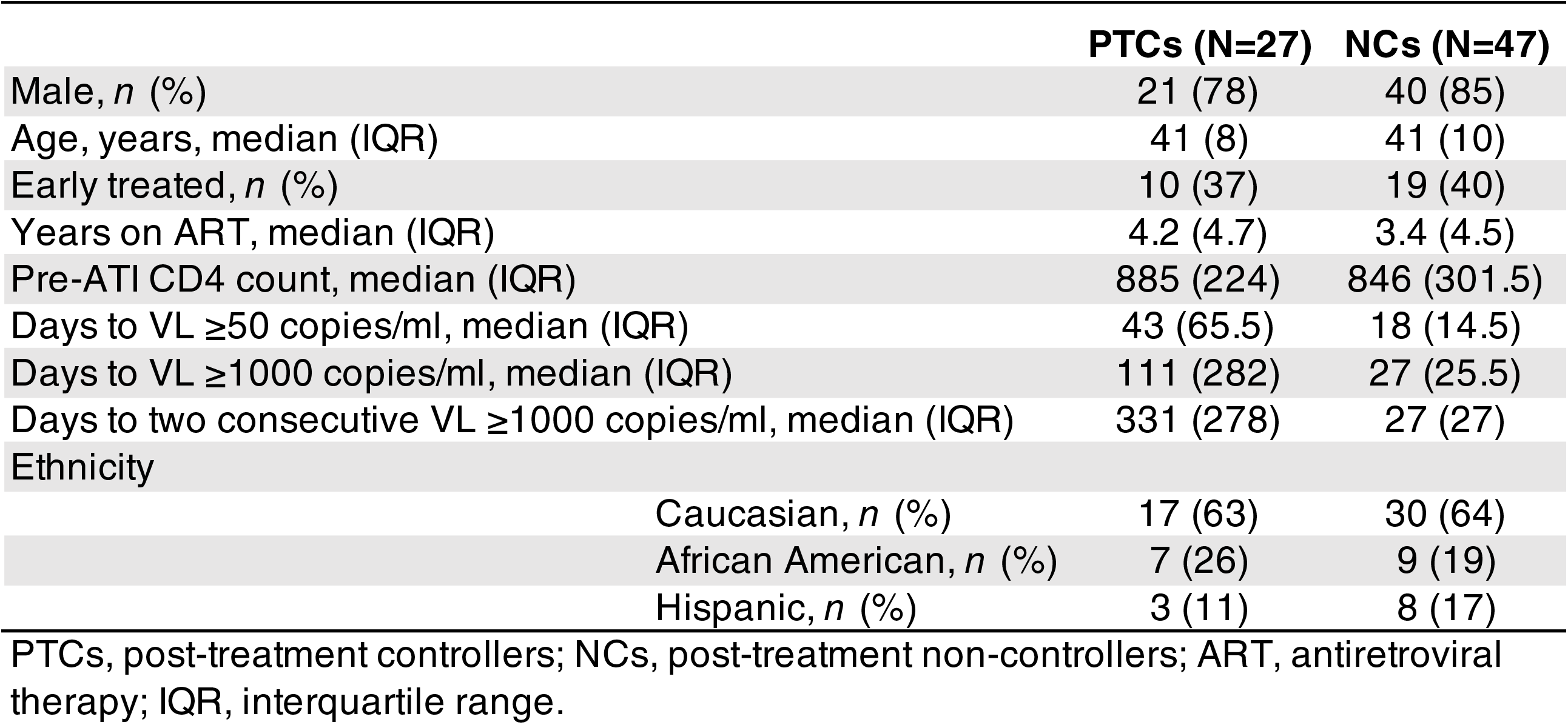
Demographic and clinical characteristics of PTCs and NCs from the ACTG cohort.

|  | **PTCs (N=27)** | **NCs (N=47)** |
| --- | --- | --- |
| Male, *n* (%) | 21 (78) | 40 (85) |
| Age, years, median (IQR) | 41 (8) | 41 (10) |
| Early treated, *n* (%) | 10 (37) | 19 (40) |
| Years on ART, median (IQR) | 4.2 (4.7) | 3.4 (4.5) |
| Pre-ATI CD4 count, median (IQR) | 885 (224) | 846 (301.5) |
| Days to VL ≥50 copies/ml, median (IQR) | 43 (65.5) | 18 (14.5) |
| Days to VL ≥1000 copies/ml, median (IQR) | 111 (282) | 27 (25.5) |
| Days to two consecutive VL ≥1000 copies/ml, median (IQR) | 331 (278) | 27 (27) |
| Ethnicity |  |  |
| Caucasian, *n* (%) | 17 (63) | 30 (64) |
| African American, *n* (%) | 7 (26) | 9 (19) |
| Hispanic, *n* (%) | 3 (11) | 8 (17) |

PTCs, post-treatment controllers; NCs, post-treatment non-controllers; ART, antiretroviral therapy; IQR, interquartile range.

**Figure 1.**
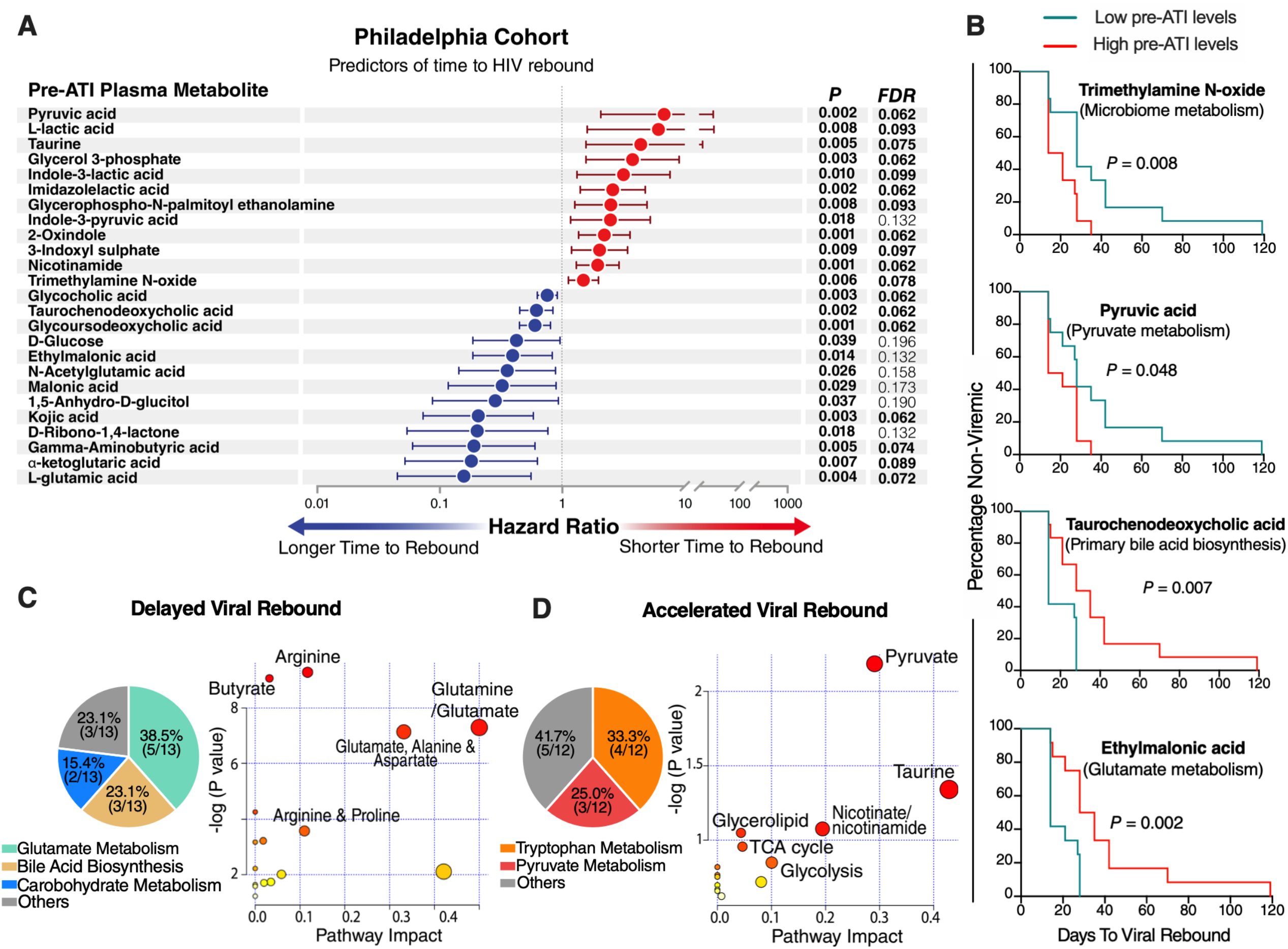
Plasma metabolites associate with time-to-viral-rebound in the Philadelphia Cohort. (**A**) Cox proportional-hazards model of metabolites that associate with longer (blue) or shorter (red) time-to-viral rebound during ATI. False Discovery Rate (FDR) was calculated using Benjamini-Hochberg correction. (**B**) Mantel-Cox test analysis of four selected metabolites from (A). Low pre-ATI levels = lower than group median; High pre-ATI levels = higher than group median. (**C**) Pathway analysis of the 13 metabolites (blue circles in A) whose pre-ATI levels associated with delayed viral rebound. Left image: multi-analysis approach combining KEGG and STRING Interaction Network. Right image: unbiased analysis using MetaboAnalyst 3.0 (http://www.metaboanalyst.ca/) where the node color is based on p-value, and the node radius is based on the pathway impact value. The pathway impact is determined by normalizing the sum of matched metabolites to the sum of all metabolites in each pathway. (**D**) Pathway analysis of the 12 metabolites (red circles in A) whose pre-ATI levels associated with accelerated viral rebound. Analysis was performed as in panel (C).

### Elevated pre-ATI levels of plasma markers of glutamate and bile acid metabolism associate with delayed viral rebound in the Philadelphia Cohort

We first aimed to examine the utility of plasma metabolites as biomarkers of time-to-HIV-rebound after ART-cessation. Towards this goal, we measured levels of plasma metabolites from the Philadelphia cohort.^35,36^ Using an untargeted mass spectrometry (MS)-based metabolomics analysis, we identified a total of 179 metabolites in plasma samples collected immediately before the ATI. Then, we applied the Cox proportional-hazards model to identify metabolomic signatures of time-to-viral-rebound. As shown in **Figure 1A**, higher pre-ATI levels of 13 plasma metabolites were significantly associated with a longer time-to-viral-rebound with *P*<0.05 and false discovery rate (FDR) <20%. In contrast, higher pre-ATI levels of 12 plasma metabolites were significantly associated with a shorter time-to-viral-rebound. When participants were separated into low or high groups by the median of each of these 25 metabolic markers, pre-ATI levels of 20 of 25 metabolites significantly indicated hazards of viral-rebound over time using the Mantel-Cox test (**Figure 1B and Supplementary Table 2**).

We next sought to determine if the 25 metabolites associated with time-to-viral-rebound shared similar metabolic pathways. Multi-analysis combining KEGG and the STRING Interaction Network (focusing on metabolite-associated enzymatic interactions) revealed that most of the 13 metabolites whose pre-ATI levels associated with a longer time-to-viral-rebound belong to two major metabolic pathways. Specifically, five metabolites lay within the anti-inflammatory glutamate/tricarboxylic acid (TCA) cycle pathway, and three were intermediates within the primary bile acid biosynthesis pathway (**Figure 1C**). Confirmatory analysis on these 13 metabolites using the MetaboAnalyst 3.0 pathway feature (http://www.metaboanalyst.ca/) showed enrichment in glutamate metabolism (*P* = 0.00068) and the bile acid biosynthesis pathway (*P* = 0.0399) (**Figure 1C and Supplementary Table 3**).

### Elevated pre-ATI levels of plasma markers of pyruvate and tryptophan metabolism associate with accelerated viral rebound in the Philadelphia Cohort

Multi-analysis of the 12 metabolites whose pre-ATI levels associated with shorter time-to-viral-rebound showed four intermediates in the tryptophan metabolism pathway and three that are central players in the pro-inflammatory pyruvate pathway (**Figure 1D**). These observations were confirmed for the 12 metabolites using MetaboAnalyst 3.0, which demonstrated enrichment in pyruvate metabolism (*P* = 0.0065) (**Figure 1D and Supplementary Table 3**). The roles of key discovered metabolites within the glutamate, bile acids, tryptophan, and pyruvate pathways are graphically illustrated in **Supplementary Figure 2**. These data reveal a previously undiscovered class of plasma metabolic biomarkers that are associated with time-to-viral rebound post-ATI. They further demonstrate that these biomarkers belong to a specific set of metabolic pathways that may play a previously unrecognized role in HIV control.

**Figure 2.**
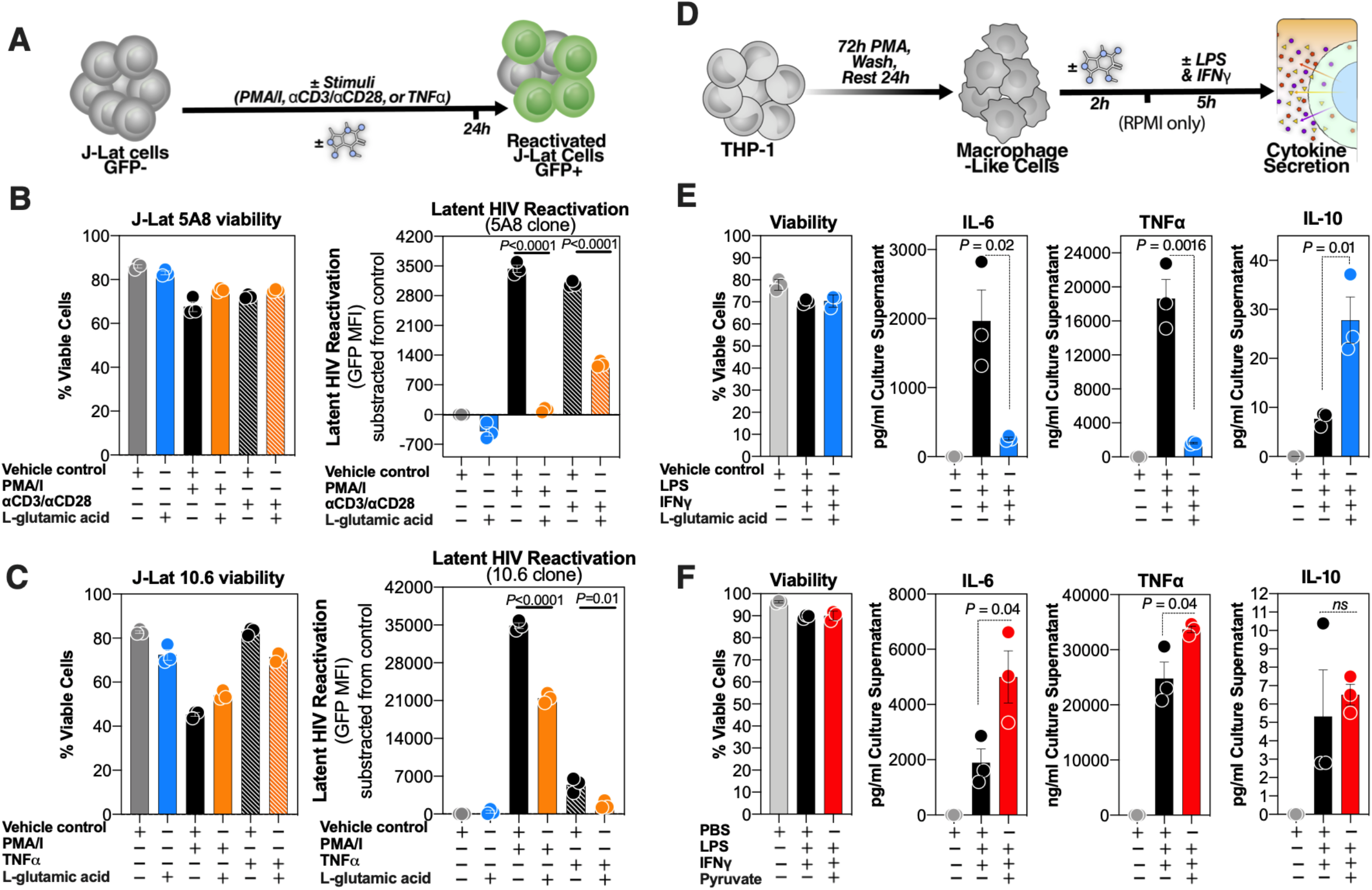
L-glutamic acid and pyruvate directly impact latent HIV reactivation and/or macrophage inflammation. (**A**) JLat 5A8 or 10.6 clones were stimulated with appropriate stimuli in the presence or absence of L-glutamic acid or vehicle control (cell culture suitable HCl solution). Geometric mean fluorescence intensity (MFI) of HIV-regulated GFP expression was measured by flow cytometry. Cell viability was determined by LIVE/DEAD aqua staining. (**B**) J-Lat 5A8 cells (n=3) and (**C**) J-Lat 10.6 cells (n=3), were treated with PMA/I (2 nM/500 nM), ImmunoCult Human CD3/CD28 T Cell Activator (25µl per 10^6^ cells), or TNFα (10ng/ml) in the presence or absence of L-glutamic acid (5mM) or appropriate control. Bar graphs display mean±SD values, and statistical comparisons were performed using two-tailed unpaired t-tests. (**D**) THP-1 cells were differentiated into macrophage-like cells using PMA. Cells were then treated with L-glutamic acid (5mM), pyruvate (2mM), or appropriate controls for 2 hours prior to LPS/IFNγ stimulation for 5 hours. Cell viability was determined by LIVE/DEAD aqua staining and cytokine secretion was measured in the supernatants using ELISA and MSD platform multiplex assay (**E**) L-glutamic acid significantly inhibited LPS/IFNγ-mediated secretion of pro-inflammatory cytokines such as IL-6 and TNFα but significantly increased the ant-inflammatory IL-10 release. Bar graphs display mean±SD, and statistical comparisons were performed using two-tailed unpaired t-tests. (**F**) Pyruvate significantly increased LPS/IFNγ-mediated secretion of IL-6 and TNFα. Bar graphs display mean±SD, and statistical comparisons were performed using two-tailed unpaired t-tests.

### L-glutamic acid and pyruvate modulate latent HIV reactivation and/or macrophage inflammation in vitro

Among the top candidate metabolic biomarkers from **Figure 1** are L-glutamic acid (glutamate metabolism) and pyruvic acid (pyruvate metabolism). The higher pre-ATI levels of L-glutamic acid and pyruvic acid associated with longer or shorter time-to-viral-rebound, respectively. These two metabolites can impact inflammation in opposing directions. Glutamate controls the anti-inflammatory TCA cycle through its conversion by glutamate dehydrogenase to α-ketoglutarate,^46,47^ whereas pyruvate is centrally positioned within the pro-inflammatory glycolytic pathway.^48-50^ We therefore sought to determine if these two metabolites exhibited a direct functional impact on latent HIV transcription and/or myeloid inflammation. We first assessed the impact of these two metabolites on latent HIV reactivation using the established “J-Lat” model of HIV latency. J-Lat cells harbor a latent, transcriptionally competent HIV provirus that encodes green fluorescent protein as an indicator of reactivation (**Figure 2A**).^51,52^ There are several clones of the J-Lat model with different characteristics, including the type of stimulation to which they respond. For example, the 5A8 is the only J-Lat clone responsive to αCD3/αCD28 stimulation. We examined the impact of L-glutamic acid and pyruvate on two J-Lat clones (5A8 and 10.6). Whereas pyruvate had no observable effect on latent reactivation for either clone (data not shown), L-glutamic acid significantly inhibited the ability of phorbol-12-myristate-13-acetate (PMA)/ionomycin or αCD3/αCD28 to reactivate latent HIV in clone 5A8 without impacting viability compared to stimuli alone controls (**Figure 2B**). L-glutamic acid also inhibited the ability of PMA/ionomycin or TNFα to reactivate latent HIV in clone 10.6 without impacting viability compared to stimuli alone controls (**Figure 2C**). These data demonstrate that a plasma metabolite, L-glutamic acid, can inhibit latent viral reactivation, consistent with the observation that pre-ATI levels of L-glutamic acid predicted a longer time-to-viral-rebound.

Beyond direct impact on latent viral reactivation, plasma metabolites may exert effects on myeloid inflammation, and such effects may underlie HIV control during ATI. This possibility was tested by examining the effects of L-glutamic acid and pyruvate on lipopolysaccharides (LPS)-mediated secretion of pro-inflammatory cytokines from THP-1 derived macrophage-like cells. These cells characterized by high basal glycolytic activity closely reflect the Warburg-like phenotype observed in HIV infected individuals,^53^ and exhibit similar inflammatory responses to primary cells under similar *in vitro* conditions.^48^ Cells were treated with L-glutamic acid, pyruvate, or appropriate controls for 2 hours before stimulating with LPS and IFNγ for 5 hours (**Figure 2D**). L-glutamic acid inhibited LPS/IFNγ-mediated production of pro-inflammatory cytokines such as IL-6 and TNFα (**Figure 2E**; other cytokines are shown in **Supplementary Figure 3A**). Consistently, L-glutamic acid also increased anti-inflammatory IL-10 secretion (**Figure 2E**). Conversely, pyruvate increased IL-6 and TNFα secretion (**Figure 2F**; other significantly regulated cytokines are shown in **Supplementary Figure 3B**). These data demonstrate that not only do some metabolites associate with time-to-viral-rebound, but also that there is a plausible, functionally significant link between these biomarkers and viral control during and following ATI.

**Figure 3.**
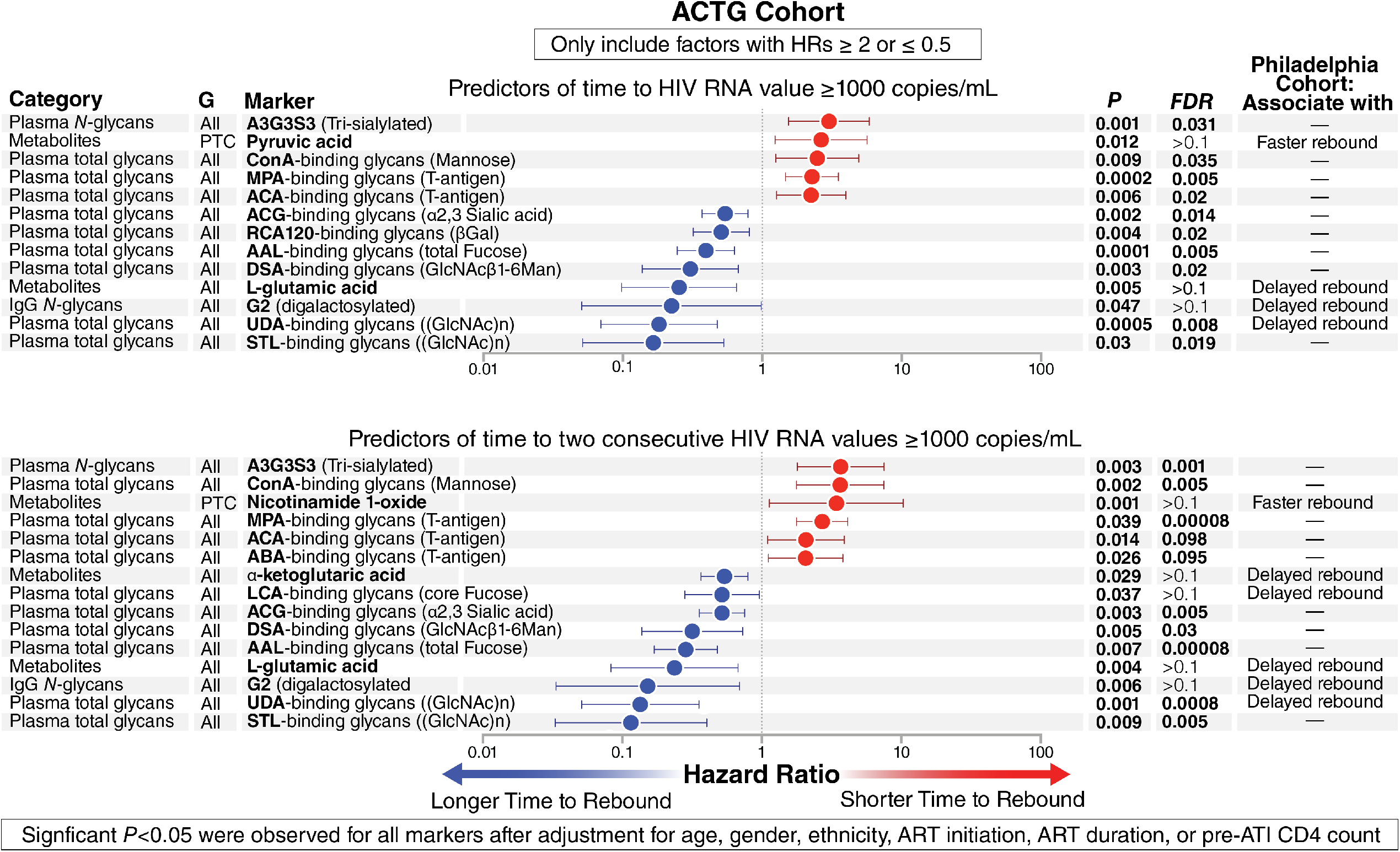
Hazard ratios of plasma glycomic and metabolic markers that associated with time-to-viral-rebound in the ACTG Cohort. Cox proportional-hazards model of glycomic and metabolic markers of time to (top panel) VL ≥1000 or (bottom panel) two constitutive VL ≥1000 within the ACTG Cohort. G = group (All = using data from all 74 participants and PTC = using data from only the 27 PTCs within the ATG Cohort). False Discovery Rate (FDR) was calculated using Benjamini-Hochberg correction.

### Pre-ATI plasma glycomic and metabolic biomarkers associate with time-to-viral-rebound in the ACTG Cohort

Our recent pilot study showed that pre-ATI levels of a specific set of glycans predicted a longer time-to-viral rebound after ART discontinuation.^35^ However, this small pilot study did not correct for confounders such as age, gender, and nadir CD4 count on viral rebound. We hypothesized that a set of plasma glycans and metabolites we identified in that pilot study,^35^ as well as in the results shown in **Figure 1**, can predict time-to-viral-rebound and/or probability-of-viral-rebound using plasma samples from a larger validation cohort, even after adjusting for potential demographic and clinical confounders. For this analysis we analyzed samples from the ACTG cohort.

We analyzed the plasma metabolome of samples collected from this cohort before ATI. A total of 226 metabolites were identified using MS-based metabolomics analysis. In addition, we applied two different glycomic technologies to analyze the plasma glycome of the same samples. First, we used capillary electrophoresis to identify the *N*-linked glycans of total plasma glycoproteins (identified 24 glycan structures, their names and structures are listed in **Supplementary Figure 4**) and isolated plasma IgG (identified 22 glycan structures, their names and structures are listed in **Supplementary Figure 5**). Second, we used a 45-plex lectin microarray to identify total (*N* and *O* linked) glycans on plasma glycoproteins. The lectin microarray enables sensitive identification of multiple glycan structures by employing a panel of 45 immobilized lectins (glycan-binding proteins) with known glycan-binding specificity, resulting in a “glycan signature” for each sample (the 45 lectins and their glycan-binding specificities are listed in **Supplementary Table 4**.).^54^

**Figure 4.**
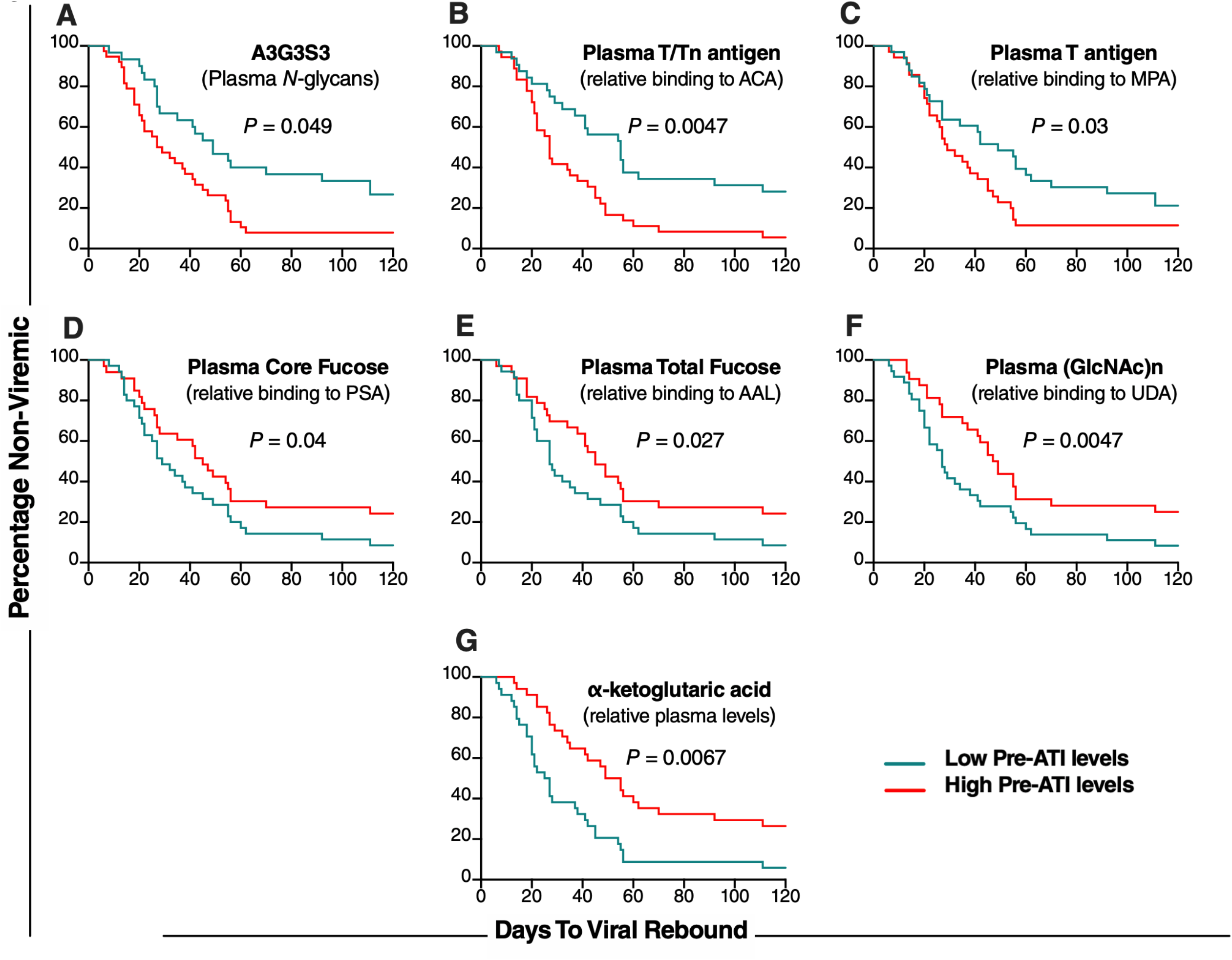
Mantel-Cox plots of plasma glycomic and metabolic markers that associated with time-to-viral-rebound in the ACTG Cohort. Graphic representation of Mantel-Cox test illustrating six glycans (A-F) and one metabolite (G) that predicted time-to-viral-rebound in Figure 3. Low pre-ATI levels = lower than the median; high pre-ATI levels = higher than the median.

**Figure 5.**
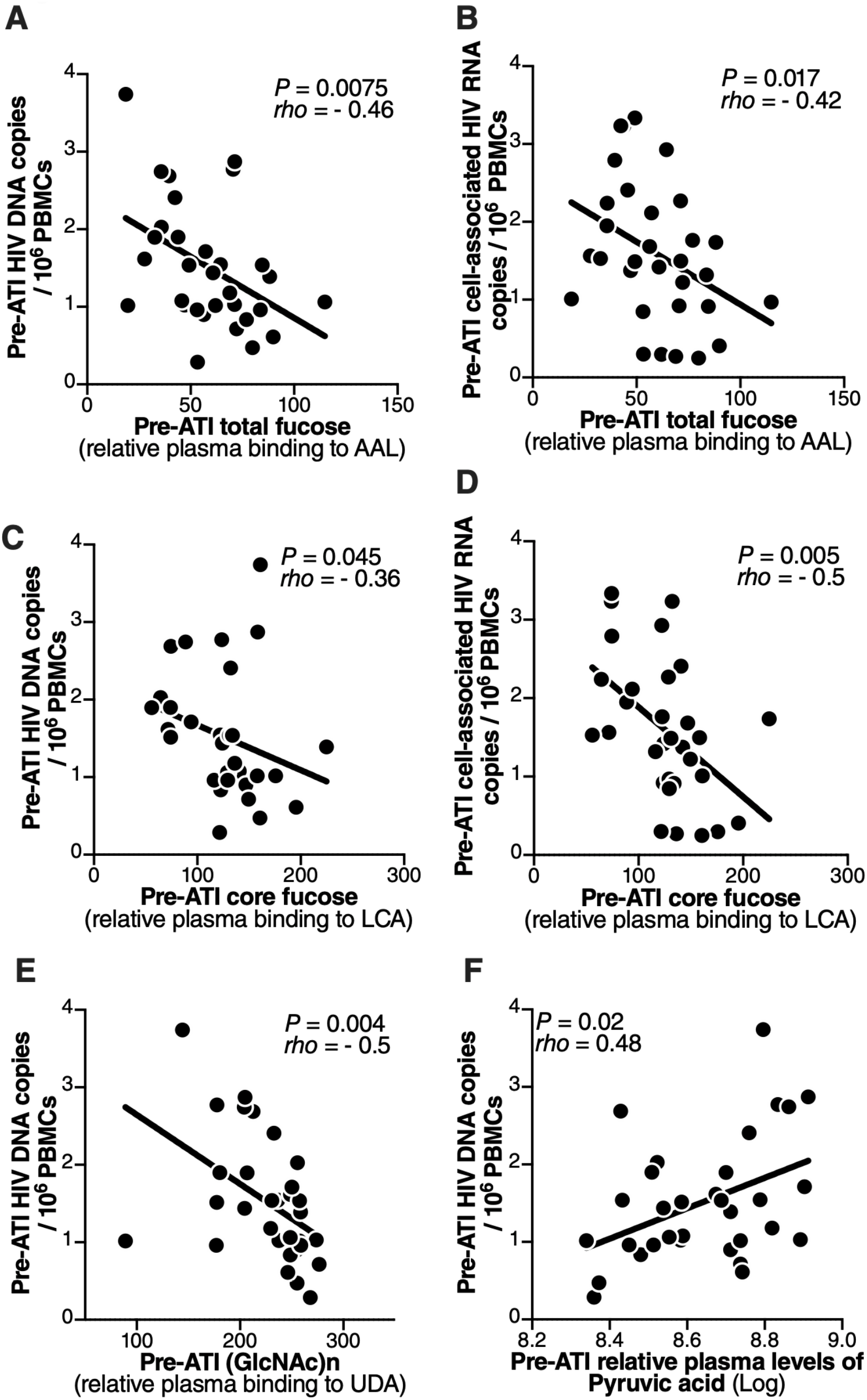
Plasma glycomic and metabolic markers of time-to-viral-rebound associate with levels of PBMC-associated HIV DNA and RNA in the ACTG Cohort. (**A-D**) Inverse associations between pre-ATI plasma levels of total fucose and core fucose, and levels of pre-ATI cell-associated (**A**,**C**) HIV DNA or (**B**,**D**) HIV RNA. (**E**,**F**) Associations between pre-ATI plasma levels of the (**E**) glycan structure (GlcNac)n or (**F**) the metabolite pyruvic acid and levels of cell-associated HIV DNA. All correlations were done using Spearman’s rank correlation coefficient tests.

We used the Cox proportional-hazards model and a set of highly stringent criteria to identify sets of glycans or metabolites whose pre-ATI levels associated with either time to VL ≥1000 (**Figure 3 top panel**) or time to two consecutive VL ≥1000 (**Figure 3 bottom panel**). To ensure high stringency, we only considered markers with a hazard ratio (HR) ≥ 2 or ≤0.5. We also included in these sets only those glycomic and metabolic markers with either FDR<0.1 or markers that emerged from the Philadelphia cohort (**Figure 1** and our previous pilot study^35^). Importantly, we only included markers that remained significant (*P*<0.05) after adjusting for age, gender, ethnicity, ART initiation (during early or chronic HIV infection), ART duration, or pre-ATI CD4 count (**Supplementary Table 5**). These combined strict criteria identified a signature that predicted shorter time-to-rebound to VL≥1000, comprising four glycan structures and one metabolite (**Figure 3 top panel, red**).These five markers include the highly sialylated plasma *N*-glycan structure (A3G3S3), GalNAc-containing glycans (also known as T-antigen; measured by binding to both MPA and ACA lectins) and the metabolite pyruvic acid. We also identified a signature that associated with a longer time-to-rebound to VL≥1000, comprising seven glycan structures and one metabolite, notably the digalactosylated G2 glycan structure on plasma bulk IgG, fucosylated glycans in plasma (binding to AAL lectin), GlcNac glycans in plasma (binding to DSA, UDA, and STL lectins), and the metabolite L-glutamic acid (**Figure 3 top panel, blue**).

Turning to markers that associated with time to two consecutive VL ≥1000, and applying the same strict criteria, we identified five glycomic markers and one metabolite whose pre-ATI levels associate with shorter time-to-rebound post-ATI, including A3G3S3 in plasma, T/Tn-antigens (binding to MPA, ACA, and ABA lectins), and the metabolite Nicotinamide (**Figure 3 bottom panel, red**). We also identified seven glycan structures and two metabolites whose pre-ATI levels predicted a longer time-to-rebound, including, G2 glycan structure on bulk IgG, core fucosylated glycans (binding to LCA lectin) in plasma, total fucosylated glycans (binding to AAL lectin) in plasma, GlcNac glycans (binding to DSA, UDA, and STL lectins) in plasma, and the metabolites oxoglutaric acid (α-ketoglutaric acid) and L-glutamic acid (**Figure 3 bottom panel, blue**). The significance of several of these markers was also confirmed using the Mantel-Cox test in an independent analysis (**Figure 4**). In sum, using stringent analysis criteria that also took into account potential confounders, we identified and validated plasma glycomic/metabolomic signatures of time-to-viral-rebound after ART discontinuation in this independent heterogeneous cohort of individuals who underwent ATI and received or not several different interventions before ATI.

### Levels of pre-ATI plasma glycomic and metabolic markers that associate with time-to-viral-rebound are linked to levels of cell-associated HIV DNA and RNA

We next examined whether the plasma glycans and metabolites (**Figure 3**) that associated with time-to-viral-rebound also reflected levels of virological markers of HIV persistence (levels of peripheral blood mononuclear cell (PBMC)-associated total HIV DNA and HIV RNA) in blood. We found that pre-ATI levels of total fucose (binding to AAL lectin), which predicted delayed viral rebound, showed a significant inverse correlation with pre-ATI levels of cell-associated HIV DNA and RNA (**Figure 5A-B**). Similarly, pre-ATI levels of core fucose (binding to LCA lectin), which also predicted delayed viral rebound, also showed an inverse correlation with pre-ATI levels of cell-associated HIV DNA and RNA (**Figure 5C-D**). Furthermore, total levels of (GlcNAc)n (binding to UDA lectin), which predicted delayed viral rebound, had an inverse correlation with levels of total HIV DNA (**Figure 5E**). Noteworthy, levels of pyruvic acid, whose pre-ATI levels predicted accelerated viral rebound, had a significant positive correlation with pre-ATI levels of cell-associated HIV DNA (**Figure 5F**). These data provide more support for a plausible mechanistic connection between our discovered plasma markers and HIV control during ATI.

### Multivariable Cox model, using Lasso technique with the cross-validation (CV), selected variables that their combination predicts time-to-viral-rebound

As a single marker would be highly unlikely to strongly predict these complex virological milestones, we next sought to apply a machine learning algorithm to identify a smaller set of plasma biomarkers (from Figure 3) that together can predict either time to VL ≥1000 or time to two consecutive VL ≥1000 better than any of these biomarkers individually. The analysis considered biomarkers, both metabolites and/or glycan structures, that emerged as significant from the ACTG cohort (**Figure 3**) and using samples with complete data set (n=70; four samples did not have a complete dataset). The machine learning algorithm, Lasso (least absolute shrinkage and selection operator) regularization, selected seven markers from among the 13 that associated with time to VL ≥1000 (**Figure 3 top panel**), whose predictive values are independent and combining them together would enhance the predictive ability of the signature compared to each of these marker alone (**Supplementary Table 6**). Indeed, a multivariable Cox regression model using these seven variables showed a concordance index (C-index) value of 74% (95% confidence interval: 68%-80%), which is significantly higher than the C-index values obtained from Cox models using each variable individually (*P*<0.05; **Supplementary Table 6**). Notably these seven markers included four whose pre-ATI levels associated with accelerated rebound, A3G3S3, T-antigen (MPA and ACA lectins binding), and the metabolite pyruvic acid. The other three markers associated with delayed rebound: total fucose (AAL lectin binding), (GlcNAc)n (STL lectin binding), and the metabolite L-glutamic acid (**Supplementary Table 6**).

Examining markers associated with time to two consecutive VL ≥1000, Lasso selected 12 markers from the 15 identified (**Figure 3 bottom panel**) whose predictive values are independent and whose combination enhanced the predictive ability of the signature compared to any single marker alone (**Supplementary Table 7**). A multivariable Cox regression model using these 12 variables showed a concordance index (C-index) value of 76.4% (95% confidence interval: 70%-84.2%), which is significantly higher than the C-index values obtained from Cox models using each variable individually (*P*<0.05; **Supplementary Table 7**). The 12 markers included some whose pre-ATI levels associated with accelerated rebound, including A3G3S3 glycans and T-antigen (ABA and ACA lectins binding) and some whose pre-ATI levels associated with delayed viral rebound, including G2 glycans, total fucose (AAL lectin binding), (GlcNAc)n (STL lectin binding), and the metabolite L-glutamic acid. (**Supplementary Table 7**). Together, these data suggest that these multivariable models of combined plasma glycans and metabolites markers warrant further exploration for their capacity to predict time-to-viral rebound in different settings.

### Pre-ATI plasma glycomic and metabolic markers distinguish post-treatment controllers (PTC) from non-controllers (NCs)

Examining the glycan structures and metabolites obtained from the ACTG cohort, we identified eight glycan structures whose pre-ATI levels were significantly different between PTCs and NCs with FDR<0.1 (**Figure 6A-H**). Among these eight glycans structures, three exhibited lower levels in the plasma of PTCs compared to NCs (FDR<0.02), including the di-sialylated glycans, A2, in total IgG glycans; the highly-sialylated glycans, A3G3S3, in plasma *N*-glycans; and T-antigen (binding to ABA lectin) (**Figure 6A-C**); and five glycans were higher in PTCs compared to NCs (FDR≤0.035). These included total fucose (binding to AAL lectin), core fucose (binding to LCA and PSA lectins), and (GlcNac)n (binding to STL and UDA lectins (**Figure 6D-H)**.

**Figure 6.**
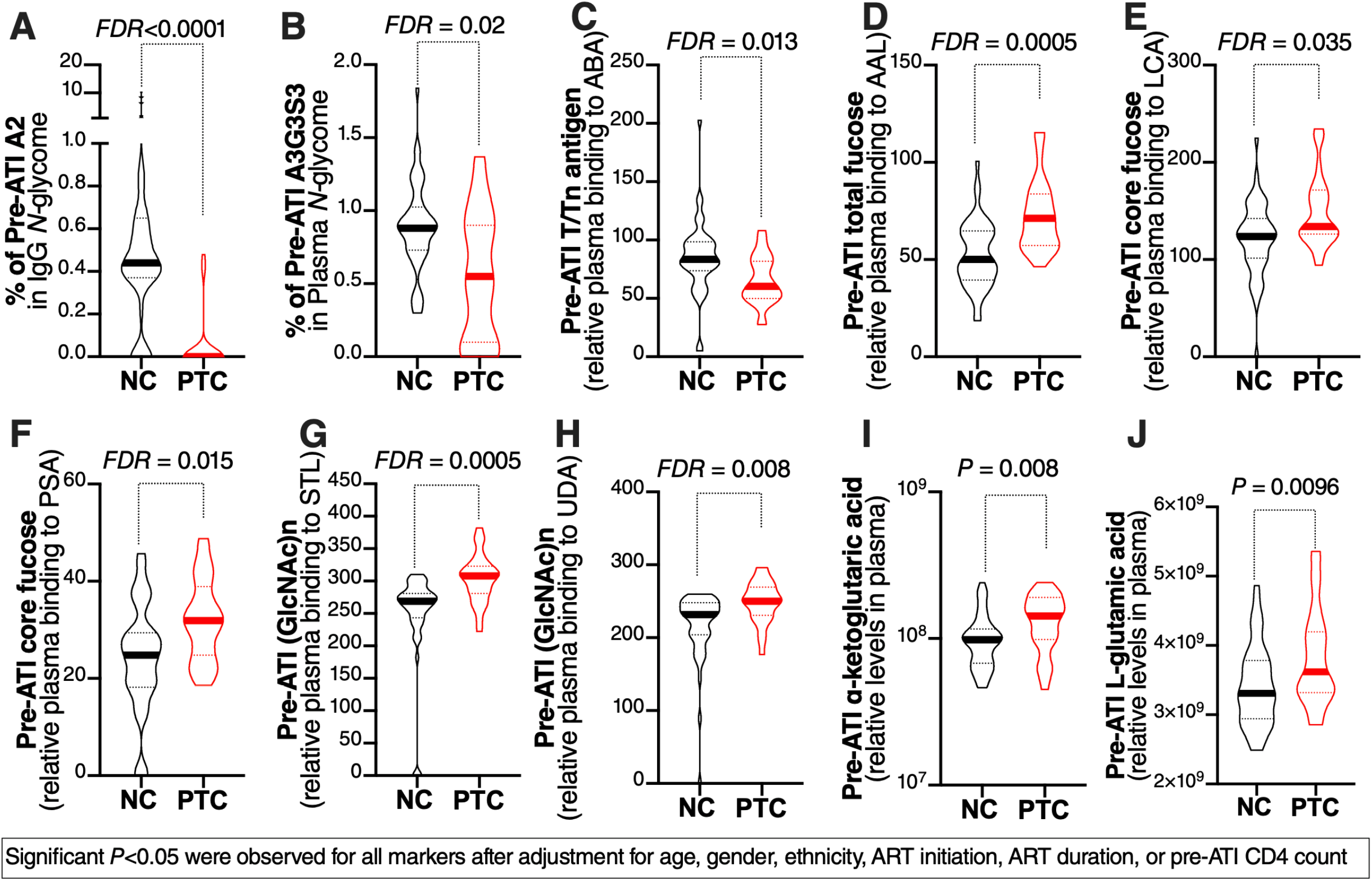
Plasma glycomic and metabolic markers that distinguish post-treatment controllers (PTCs) from non-controllers (NCs). Pre-ATI levels of three glycan structures were are lower in PTCs compared to NCs: (**A**) the disialylated glycans, A2, in the IgG glycome, (**B**) the highly sialylated glycan structure (A3G3S3), and (**C**) T/Tn antigen (measured as binding to ABA lectin). Pre-ATI levels of four glycan structures were higher in PTCs compared to NCs: (**D**) total fucose (binding to AAL lectin) in plasma, (**E**-**F**) core fucose (binding to LCA and PSA lectins) in plasma, and (**G-H**) (GlcNAc)n (binding to STL and UDA lectins). Pre-ATI levels of two metabolites were higher in PTCs compared to NCs: (**I**) α-ketoglutaric acid and (**J**) L-Glutamic acid. All statistical comparisons were performed using a Mann-Whitney test. Truncated violin plots showing median. False Discovery Rate (FDR) was calculated using Benjamini-Hochberg (BH) correction.

Examining metabolites, we found that pre-ATI levels of α-ketoglutaric acid and L-glutamic acid, both of which predicted delayed viral rebound, were higher in the plasma of PTCs compared to NCs (*P*<0.01, **Figure 6J-I**). Importantly, for both glycans and metabolites, we only selected markers whose levels remained different (*P*<0.05) between PTCs and NCs after adjusting for age, gender, ethnicity, ART initiation, ART duration, or pre-ATI CD4 count (**Supplementary Table 8**). Together, these data suggest that a selective set of plasma glycans and metabolites can distinguish PTCs from NCs and may be used to predict the probability of viral rebound (i.e., the likelihood of PTC phenotype after ATI).

### Multivariable logistic model, using CV Lasso technique, selected variables that their combination predicts risk of viral rebound

We next applied the Lasso regularization to select from among the ten markers in Figure 6 a set of markers whose combined predictive utility is better than the predictive utility of any of these 10 markers individually. The analysis used biomarkers that emerged as significant from the ACTG cohort (**Figure 6**) and using samples with complete data set (n=70). Lasso selected seven markers from the 10 identified as able to distinguish PTCs from NCs (**Figure 6**) that their predictive values are independent and combing them together would enhance the predictive ability of the signature compared to each of these markers alone (**Supplementary Table 9**). Indeed, a multivariable logistic regression model using these seven variables showed an area under the ROC curve (AUC) value of 97.5% (**Figure 7A**; 95% confidence interval: 94% -100%), which is significantly higher than the AUC values obtained from logistic models using each variable individually (*P*<0.05; **Supplementary Table 9**). These seven markers included three whose pre-ATI levels are lower in PTCs compared to NCs, namely A2, A3G3S3, and T-antigen (ABA lectin binding), and four whose pre-ATI levels were higher in PTCs compared to NCs, namely total fucose (AAL lectin binding), core fucose (LCA lectin binding), (GlcNAc)n (STL lectin binding), and the metabolite L-glutamic acid (**Supplementary Table 9**).

**Figure 7.**
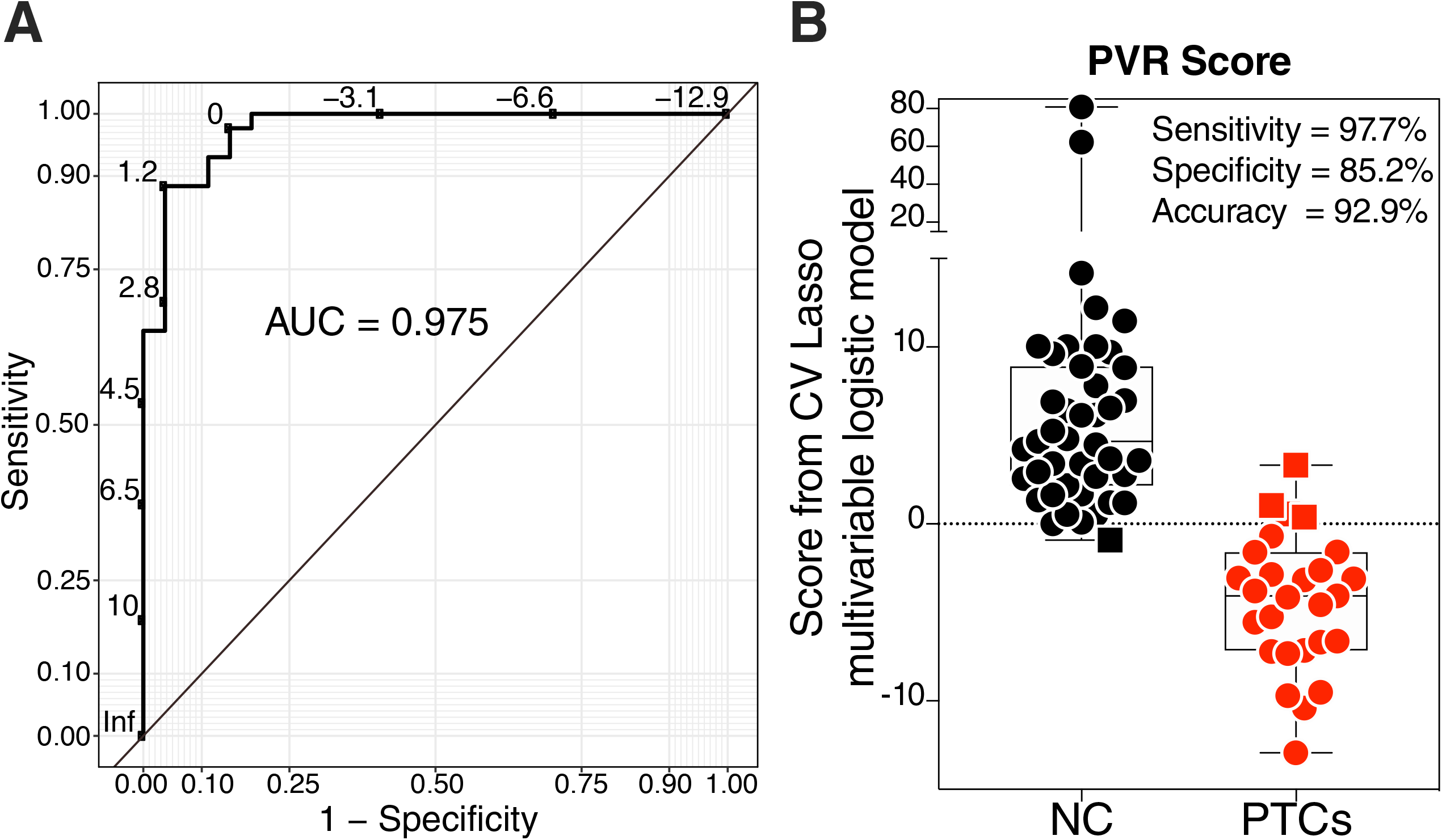
A multivariable logistic model using Lasso selected variables predicts probability of viral remission post ATI. The machine learning algorithm, Lasso (least absolute shrinkage and selection operator) regularization, selected seven markers from the 10 markers in Figure 6. Analysis using this model demonstrates that when these seven markers are combined, their predictive ability is better than the predictive ability of any marker individually (Supplementary Table 9). (**A**) Receiver operator characteristic (ROC) curve showing the area under the curve (AUC) is 97.5% from the multivariable logistic regression model with seven variables. (**B**) Coefficients from the multivariable logistic model were used to estimate risk score for each individual and then tested for the ability of these scores to accurately classify PTCs and NCs at an optimal cut-point. The model was able to correctly classify 97.7 of NCs (sensitivity), 85.2% of PTCs (specificity) with overall accuracy of 92.9%. Squares represent individuals the model failed to identify correctly.

Next, a risk score predicting NC was estimated for each individual using the multivariable logistic model. We then examined the ability of these risk scores to classify PTCs and NCs from the ACTG cohort. As shown in **Figure 7B**, the model was able to correctly classify 97.7% of NCs (sensitivity), and 85.2% of PTCs (specificity) with overall accuracy of 92.9%. This analysis highlights the potential utility of this risk score estimated from the multivariable model combining six plasma glycans and one metabolite, to predict the risk of NC post-ATI. This prediction can be utilized to select individuals likely to achieve PTC phenotype during HIV cure-focused clinical trials, to be included in ATI studies. In addition, the markers that are included in this model might also serve as windows into the mechanisms that contribute to the PTC phenotype.

## DISCUSSION

In this study we identified and initially validated pre-ATI plasma glycomic and metabolomic biomarkers of both duration and probability of viral remission after treatment interruption. We observed a significant overlap between plasma markers that predicted time-to-viral rebound and markers that predicted probability of viral rebound (i.e., predicted the PTC phenotype in comparison to the NC phenotype). Specifically, pre-ATI plasma levels of the anti-inflammatory L-glutamic acid, *N*-Acetylglucosamine (GlcNac), and fucose were associated with both delayed rebound and higher likelihood to achieve viral remission. Whereas pre-ATI plasma levels of the highly-sialylated A3G3S3 and GalNAc-containing glycans (T/Tn-antigens) were associated with both accelerated rebound and lower likelihood of achieving viral remission. Notable differences included the digalactosylated G2 glycan on IgG glycome whose pre-ATI levels associated with longer time-to-viral-rebound but not probability of viral rebound; and the di-sialylated IgG glycan, A2, whose pre-ATI levels associated with higher probability of viral rebound but not with time-to-viral rebound.

It is not surprising that a single marker cannot highly predict these complicated virological milestones (time to and probability of viral rebound). Therefore, we applied machine learning algorithms to select the smallest number of variables that, when combined, maximizes the predictive utility of our signatures. The variables selected by CV Lasso technique, when used in multivariate models, were able to predict time-to-viral rebound using Cox models with C-index of 74-76% and probability of viral rebound using logistic model with AUC of 97.5%. The utility of these multivariable models to be used in HIV cure-directed clinical trials warrants further investigation. Upon validation, these models could have a profound impact on the HIV cure field by mitigating the risk of ATI during HIV cure-focused clinical trials and provide means for selecting only the most promising therapies and most likely individuals to achieve viral remission to be tested by ATIs.

Beyond their utility as biomarkers, these metabolic and glycomic signatures of viral rebound represent an opportunity to better understand the host milieu preceding a viral rebound. The likelihood of viral rebound and viral remission after ART cessation is likely a function of both the size of the inducible replication-competent HIV reservoir and the host environment that influences inflammatory and immunological responses.^55^ The on-going efforts by many groups to understand the quantitative and qualitative nature of the HIV reservoir are critical to understanding the virological basis of viral rebound.^56-58^ However, complementary studies are also needed to understand host determinants of inflammatory and immunological states that also may impact post-treatment control of HIV. Our functional analyses on two of these biomarkers (L-glutamic acid and pyruvic acid in Figure 2) suggest that our signatures have a potential functional significance for HIV post-treatment control. These markers may directly impact latent HIV reactivation or may indirectly condition the host environment with differential levels of inflammation that might impact viral reactivation, cellular processes, and immunological functions during ATI. The potential direct and indirect functional significance of each of the key variable in our models warrants further investigations as they can serve as windows into the mechanisms that contribute to post-ART HIV control.

Our data obtained from two independent cohorts suggest that the bioactive plasma metabolites might not only predict duration and probability of viral remission, but also actively contribute to it. Our *in vivo* data showed that the pre-ATI levels of L-glutamic acid predict a delayed viral rebound and a higher probability of viral remission. Indeed, our *in vitro* validation experiments showed that L-glutamic acid can directly suppress HIV reactivation and suppress LPS and IFNγ-mediated inflammation of myeloid cells. It has been argued that L-glutamic acid, through its conversion to α-ketoglutarate, fuels the TCA cycle/oxidative phosphorylation, which is typically regarded to be an anti-inflammatory metabolic signature.^59^ TCA cycle metabolites may regulate immune processes through epigenetic modifications such as DNA methylation,^60^ which may directly impact proviral reactivation. This is consistent with our *in vivo* and *in vitro* data on L-glutamic acid. In contrast to L-glutamic acid, our *in vivo* data showed that elevated pre-ATI levels of the pro-inflammatory pyruvic acid are associated with accelerated viral rebound. We also observed a significant positive correlation between pre-ATI levels of plasma pyruvic acid and total HIV DNA, a marker for reservoir size. Our *in vitro* data confirmed these *in vivo* observations and showed that pyruvate can induce a pro-inflammatory phenotype in myeloid cells upon stimulation. Aerobic glycolysis, where pyruvate is converted into lactate, drives pro-inflammatory M1-macrophage polarization,^61^ in the context of HIV infection.^48,62^ This is consistent with our *in vivo* and *in vitro* data on pyruvate. While no studies have evaluated the impact of plasma metabolic alterations in ATI, one study observed a glycolytic plasma profile in transient HIV elite controllers (TECs) compared to persistent elite controllers (PECs).^63^ Moreover, glutamic acid was shown to be elevated in PECs compared to TECs,^63^ corresponding to our observation that glutamate metabolism associated with delayed time to HIV rebound. In totality, a global Warburg phenotype has now emerged as a classic manifestation of HIV infection.^53,64,65^ Thus, the plasma metabolite signatures we observed are likely a snapshot of the global and intrinsic cellular metabolic flux that occurs during ATI in individual patients.

In addition to L-glutamic acid and pyruvic acid, other intriguing plasma metabolites emerged from the analysis of the Philadelphia cohort. Among the plasma markers associated with delayed viral rebound is ethylmalonic acid. Ethylmalonic acid is central in the metabolism of butyrate, a short-chain fatty acid produced by the gut microbiota and known for its anti-inflammatory effects.^66^ Another group of metabolites, consisting of indole-3-pyruvic acid, indole-3-lactic acid, 3-indoxyl sulphate, and 2-oxindole, characterized accelerated rebound and may reflect a biochemical manifestation of dysbiosis of gut bacteria resulting in tryptophan catabolism.^67^ Indeed the tryptophan metabolic pathway was highlighted as one of the main metabolic pathways associated with accelerated viral rebound. Although it was not mechanistically interrogated, a positive association between plasma indoleamine 2,3-dioxygenase (IDO) activity (an immunoregulatory enzyme that metabolizes tryptophan) and total HIV DNA in peripheral blood has been established.^68^ Impaired intestinal barrier integrity is a classical feature of HIV infection, characterized by dysbiosis and increased microbial by-products that drive systemic and mucosal inflammation.^69,70^ Microbes with the capacity to catabolize tryptophan have been linked to adverse HIV disease progression,^71^ at least in part due to induction of IDO1 that interferes with Th17/Treg balance in the periphery and gut.^72^ Our data highlight previously unrecognized interactions between the gut microbiome, its metabolic activity, and HIV persistence. Understanding these potential multi-nodal complex relationships during ART and post ATI warrants further investigation.

Similar to metabolites, glycans on glycoproteins are bioactive molecules and can play significant roles in mediating immunological functions. For example, antibody glycans can alter an antibody’s Fc-mediated innate immune functions, including ADCC and several pro- and anti-inflammatory activities.^22-25^ Among glycans on antibodies, the presence of core fucose results in a weaker binding to Fcγ receptor IIIA and reduces ADCC.^73^ The same occurs with terminal sialic acid, which reduces ADCC.^74-76^ On the other hand, terminal galactose induces ADCC.^77^ In three independent geographically-distinct cohorts, two studied in our previous plot study^35^ and the ACTG cohort studied in the current study, we observed a significant association between pre-ATI levels of the digalactosylated non-fucosylated non-sialylated glycan, G2, and delayed viral rebound. G2 is the only IgG glycan trait that is terminally galactosylated, non-fucosylated, and non-sialylated (**Supplementary Figure 5**) which is compatible with higher ADCC activity. Similar to our pilot study,^35^ we observed a link between plasma levels of *N*-acetyl-glucosamine (GlcNAc) and delayed viral rebound. GlcNAc has been reported to have an anti-inflammatory impact during several inflammatory diseases by modulating NFκB activity.^78-80^ Investigating the potential direct impact of these glycans on innate immune functions and inflammation, and how this affects HIV control during ART, warrants further investigations.

Glycoproteins can also be shed from cells in different organs; therefore, their characteristics can reflect these cells’ functions. Glycans on the cell surface are involved in signaling cascades controlling several cellular processes.^81-83^ It is not clear how the higher pre-ATI levels of plasma fucose, which associate with both delayed viral rebound (in our pilot study^35^ as well as the current study) and higher likelihood for PTC status post-ATI, can directly impact viral control during ATI. Nor is it clear how the higher pre-ATI levels of plasma GalNAc-containing glycans (T/Tn antigens), which associate with both accelerated viral rebound and lower likelihood for PTC status, can directly impact viral control during ATI. However, these higher levels might reflect differential levels of these glycans on cells in different organs. For example, T-antigens (tumor-associated antigen) and Tn antigen are *O*-glycans that are truncated and have incomplete glycosylation, commonly present in cancerous cells, and have been used as tumor markers.^84-87^ These GalNAc-containing glycans expressed on some normal immune cells (such as T cells) are ligands of the macrophage galactose type lectin (MGL) that is expressed on activated antigen presenting cells (APCs). MGL interacts with GalNAc-containing glycans on T cells to induce T cell dysfunction.^88^ Our data show that higher levels of these antigens in plasma ae associated with accelerated rebound and lower likelihood of viral remission and raise the question of whether these glycan levels reflect an immunosuppressive environment in NCs and those who rebound fast. Future studies are needed to examine the direct impact of these glycans on HIV control and/or the potential meaning of their levels as reflections of cellular functions in different tissues during ATI in HIV+ individuals.

We ensured the inclusion in our multivariate models of only metabolic and glycomic markers whose significance was not dependent on several demographic and clinical confounders such as age, gender, ethnicity, ART initiation during early versus chronic stages of HIV infection, duration of ART, and pre-ATI CD4 count, as all of these markers can influence HIV reservoir size and/or our metabolic/glycomic signatures. However, other potential confounders could impact our results including ART regimen, diet, co-morbidities, co-infections, and other medications. Investigating these other confounders as well as investigating geographically distinct and pediatric cohorts should be the subject of future studies. We examined the links between our glycomic and metabolic signatures and levels of cell-associated HIV DNA and RNA in the blood. However, the majority of HIV DNA and RNA harbor mutations and/or deletions, rendering them defective.^89,90^ It will be important to examine the potential links between these plasma markers and the level of intact and inducible HIV reservoirs in the blood and tissues. In addition, it will be important, in future studies, to examine the potential links between these plasma markers and host immunological and inflammatory responses. Despite these shortcomings, our study represents the first to identify a set of non-invasive, previously unrecognized class of plasma molecules (glycans and metabolites) that can be used as biomarkers of HIV remission. These signatures of viral rebound were obtained using two independent cohorts of ATI and after applying stringent criteria to avoid the potential impact of several confounders. Our machine learning algorithms also identified a combination of these markers that can enhance their predictive value. Our novel signatures, upon further validation, have the potential to fill a major gap in the HIV cure field through their usage as biomarkers of viral rebound during HIV cure-focused clinical trials. In addition, these results open new mechanistic avenues to better understand the fundamental biological processes, including carbohydrate metabolism, that may regulate HIV control during ART and post-ATI.

## METHODS

### Study cohorts

Analyses were performed from banked plasma samples of two different cohorts that underwent analytical treatment interruption (ATI): (1) Philadelphia Cohort and (2) ACTG cohort. In the Philadelphia cohort,^35,36^ 24 HIV+ individuals on suppressive ART underwent an open-ended ATI without concurrent immunomodulatory agents.^35,36^ The ACTG cohort combined 74 HIV-infected ART-suppressed participants who underwent ATI from six ACTG ATI studies (ACTG 371,^37^ A5024,^38^ A5068,^39^ A5170,^40^ A5187,^41^ and A5197).^42^ 27 of these 74 individuals exhibited a PTC phenotype post-ATI, i.e. these individuals remained off ART for ≥24 weeks post-treatment interruption, sustained virologic control for at least 24 weeks, maintained viral load (VL) ≤400 copies for at least 2/3 of time points, had plasma drug level testing performed, and had no evidence of spontaneous control pre-ART. The remaining 47 cohort members were non-controllers (NCs) who exhibited virologic rebound before meeting PTC criteria. These two groups were matched for gender, age, % treated at the early stage of HIV infection, ART duration, pre-ATI CD4 count, and ethnicity, as shown in **Table 1**. All analyses were performed on samples collected immediately before ATI in both cohorts.

### Plasma untargeted metabolomics analysis

Metabolomics analysis was performed as described previously.^91^ Briefly, polar metabolites were extracted from plasma samples with 80% methanol. A quality control (QC) sample was generated by pooling equal volumes of all samples and was injected periodically during the sequence of LC-MS runs. LC-MS was performed on a Thermo Scientific Q Exactive HF-X mass spectrometer with HESI II probe and Vanquish Horizon UHPLC system. Hydrophilic interaction liquid chromatography was performed at 0.2 ml/min on a ZIC-pHILIC column (2.1 mm × 150 mm, EMD Millipore) at 45 °C. Solvent A was 20 mM ammonium carbonate, 0.1% ammonium hydroxide, pH 9.2, and solvent B was acetonitrile. The gradient was 85% B for 2 min, 85% B to 20% B over 15 min, 20% B to 85% B over 0.1 min, and 85% B for 8.9 min. All samples were analyzed by full MS with polarity switching. The QC sample was also analyzed by data-dependent MS/MS with separate runs for positive and negative ion modes. Full MS scans were acquired at 120,000 resolution with a scan range of 65-975 m/z. Data-dependent MS/MS scans were acquired for the top 10 highest intensity ions at 15,000 resolution with an isolation width of 1.0 m/z and stepped normalized collision energy of 20-40-60. Data analysis was performed using Compound Discoverer 3.1 (ThermoFisher Scientific). Metabolites were identified by accurate mass and retention time using an in-house database generated from pure standards or by MS2 spectra using the mzCloud spectral database (mzCloud.org) and selecting the best matches with scores of 50 or greater. Metabolite quantification used peak areas from full MS runs and were corrected based on the periodic QC runs. Peak areas from samples of the ACTG study were normalized to the summed area for identified metabolites in each sample.

### In-vitro examination of the impact of L-glutamic acid on latent HIV reactivation

J-Lat cells were used as model of HIV latency. J-Lat cells harbor latent, transcriptionally-competent HIV provirus that encodes green fluorescent protein (GFP) as an indicator of viral reactivation.^51,52^ Levels of latent HIV transcription after stimulation can be measured using flow cytometry. L-glutamic acid was purchased from Sigma (catalog# 49449-100G) and was dissolved in cell-culture compatible HCl solution (Sigma catalog# H9892-100ML). J-Lat 5A8 clone was kindly provided by Dr. Warner Greene (The Gladstone Institute of Virology and Immunology). J-Lat clone 10.6 (catalog number 9849) was provided by the NIH AIDS Reagent Program (Germantown, MD). Cells from different clones of J-Lat (5A8 and 10.6) were cultured at 1×10^6^ cells/ml in cultured in R10 media (complete RPMI 1640 medium supplemented with 10% fetal bovine serum (FBS)), and were stimulated with PMA/ionomycin (16 nM/500 nM-Sigma catalog# P8139/ catalog# I0634-1MG, respectively) or ImmunoCult Human CD3/CD28 T Cell Activator (Stem cell catalog# 10971), or TNFα (10 ng/ml; Stem Cell catalog# 78068.1) in the presence of HCl solution as a control. J-Lat cells were also treated with L-glutamic acid (5mM) in the presence or absence of the above stimulators. After 24 hours, cells were stained with live/dead marker (Thermo catalog# L34966) and GFP Mean Fluorescence intensity (MFI) was measured by LSR II flow cytometer and FACSDiva software.

### In-vitro examination of the impact of L-glutamic acid and pyruvate on myeloid inflammation

THP-1 cell line (catalog number 9942) was provided by the NIH AIDS Reagent Program (Germantown, MD). THP1 cells were plated in 24-well plates at a density of 7×10^5^ cells per well. To differentiate them into macrophages-like, 100nM of PMA (Sigma catalog# P8139) was added and incubated for 72hours. After incubation, media was aspirated, and each well was gently washed twice with R10 media. Cells were then rested for 24 hours on R10 media without PMA. After 24 hours, cells were washed again with serum-free (no FBS) RPMI 1640 media and kept on this media for the rest of experiment. Macrophage-like THP1 cells were pre-incubated with L-glutamic acid (5mM) or Sodium Pyruvate solution (2mM, Sigma catalog# S8636-100ml) for 2 hours before stimulating with Escherichia coli serotype O127:B8 LPS (50ng/ml; Sigma catalog# L3129-10MG) and IFNγ (10ng/ml; R&D Systems catalog# 285-IF-100, respectively). After 5 hours of incubation with LPS/IFNγ, culture supernatants were collected for cytokine quantitation. Supernatant levels of IL-10, IL-12p70, IL-13, IL-1β, IL-2, IL-4, IL-6, and IL-8 were determined using U-PLEX Proinflam Combo 1 (Meso Scale Diagnostic # K15049k-1) according to manufacture. Levels of TNF-α were quantified using DuoSet ELISA kits (R&D Systems catalog# DY210-05).

### IgG isolation

Bulk IgG was purified from 50µl plasma using Pierce™ Protein G Spin Plate (Thermo Fisher catalog# 45204). IgG purity was confirmed by SDS gel.

### N-glycan analysis using capillary electrophoresis

For both plasma and bulk IgG, *N*-glycans were released using peptide-N-glycosidase F (PNGase F) and labeled with 8-aminopyrene-1,3,6-trisulfonic acid (APTS) using the GlycanAssure APTS Kit (Thermo Fisher cat. A33952), following the manufacturer’s protocol. Labeled *N*-glycans were analyzed using the 3500 Genetic Analyzer capillary electrophoresis system. IgG *N*-glycan samples were separated into 22 peaks and total plasma *N*-glycans into 24 peaks. Relative abundance of *N*-glycan structures was quantified by calculating the area under the curve of each glycan structure divided by the total glycans using the Applied Biosystems GlycanAssure Data Analysis Software Version 2.0.

### Glycan analysis using lectin array

To profile the plasma total glycome, we used the lectin microarray as it enables analysis of multiple glycan structures; it employs a panel of 45 immobilized lectins with known glycan-binding specificity. Plasma proteins were labeled with Cy3 and hybridized to the lectin microarray. The resulting chips were scanned for fluorescence intensity on each lectin-coated spot using an evanescent-field fluorescence scanner GlycoStation Reader (GlycoTechnica Ltd.), and data were normalized using the global normalization method.

### Quantification of HIV DNA and CA-RNA

Cell-associated (CA)-RNA and DNA were isolated from cryopreserved peripheral blood mononuclear cells (PBMCs) using the AllPrep DNA/RNA Mini Kit (Qiagen). Unspliced CA-RNA and total HIV DNA levels were quantified using a real-time PCR approach with primers/probes targeting conserved regions of HIV LTR/gag as previously described.^8,92^ The CA-RNA assay measures levels of unspliced transcripts, which are late RNA products necessary for the creation of HIV structural proteins and remains one of the most commonly used assay in HIV curative studies.^93-96^ Cell numbers were quantified by the real-time PCR measurement of CCR5 copy numbers. Cellular integrity for RNA analysis was assessed by the measurement of total extracted RNA and evaluation of the IPO-8 housekeeping gene.^97^

### Statistical analysis

For each of the studied biomarkers, data distribution was first examined, and appropriate data transformation was made for further analysis. Data from metabolic analysis and lectin array were log^2^-transformed before analysis. Two-group t-tests or Mann-Whitney tests were used to determine the difference between two groups. Spearman’s rank correlation coefficient was used to evaluate correlations. For binary outcome (NC vs. PTC) or time-to-viral-rebound, logistic or Cox regression models with or without adjusting for confounders were used to assess the association between a biomarker and outcome, respectively. False discovery rates (FDR) were calculated using Benjamini-Hochberg correction. To explore biomarkers that could be predictors of clinical outcomes, specific sets of biomarkers were identified among those with FDR<0.1. Variables for the multivariable models were selected from the identified specific sets of biomarkers using Lasso technique with the cross-validation (CV) selection option by separating data in 5-fold. Due to this exploratory study with modest sample size, variables selection was determined using 100 independent rounds runs of CV Lasso with minimum tuning parameter lambda. The biomarkers that were selected 80 or more times from 100 runs were used as final set of predictors in our models. The predictive ability of final logistic model and Cox model were assessed by AUC and C-index. GraphPad Prism 6, Stata 16, and R were used for data analysis.

### Data availability

The authors declare that data supporting the findings of this study are available within the paper and its supplementary information files. Further raw data not included in our findings are available from the corresponding author upon request.

## SUPPLEMENTARY MATERIALS

**Supplementary Figure 1**. Longitudinal viral loads of PTCs and NCs from the ACTG cohort.

**Supplementary Figure 2. Pathways enriched in the analysis of plasma metabolites for time-to-viral-rebound in the Philadelphia Cohort**. (A) Glutamate metabolism pathway. (B) Primary bile acid biosynthesis pathway. (C) Pyruvate metabolism pathway. (D) Tryptophan metabolism pathway. Metabolites in blue circles are those in which their pre-ATI levels associated with a delayed viral rebound. Metabolites in red circles are those in which their pre-ATI levels associate with accelerated viral rebound.

**Supplementary Figure 3. Effects of L-Glutamic acid and pyruvate on myeloid inflammation**. THP-1 cells were differentiated into macrophage-like cells using PMA. Macrophage-like cells were treated with L-Glutamic acid, Pyruvate, or appropriate controls for 2 hours before stimulating with LPS and IFNγ for 5 hours. (**A**) Impact of L-Glutamic acid on LPS/IFNγ mediated cytokine secretion. (**B**) Impact of Pyruvate on LPS/IFNγ mediated cytokine secretion. Mean±SD is displayed as bar charts, and statistical comparisons were performed using two-tailed unpaired t-tests.

**Supplementary Figure 4**. The structures and names of *N*-glycans identified in plasma by capillary electrophoresis.

**Supplementary Figure 5**. The structures and names of *N*-glycans identified in isolated plasma IgG by capillary electrophoresis.

**Supplementary Table 1**. Clinical and demographic data of the Philadelphia Cohort.

**Supplementary Table 2**. A list of metabolites which their pre-ATI levels associate with time-to-viral-rebound in the Philadelphia Cohort. Analysis was conducted using the Proportional Cox Hazard Model as well as the Mantel-Cox test.

**Supplementary Table 3**. Pathways enriched by metabolites associated with time-to-viral-rebound in the Philadelphia Cohort. Pathway analysis utilized the MetaboAnalyst 3.0 program (http://www.metaboanalyst.ca/).

**Supplementary Table 4**. Lectins used in the 45-plex lectin microarray and their glycan-binding specificity.

**Supplementary Table 5**. Plasma glycomic and metabolomic predictors of time-to-viral-rebound in the ACTG Cohort after adjusting for potential confounders.

**Supplementary Table 6**. Comparisons of C-index values between each univariate Cox model and the multivariable Cox model with Lasso selected variables predicting time to VL ≥1000.

**Supplementary Table 7**. Comparisons of C-index values between each univariate Cox model and the multivariable Cox model with Lasso selected variables predicting time to two consecutive VL ≥1000.

**Supplementary Table 8**. Plasma glycomic and metabolic markers that distinguished PTCs from NCs in the ACTG Cohort after adjusting for potential confounders.

**Supplementary Table 9**. Comparisons of AUC values between each logistic model employing single marker predictors versus multivariable logistic regression with Lasso selected marker set predicting PTC and NC status.

## AUTHOR CONTRIBUTIONS

M.A-M conceived and designed the study. L.B.G carried out the majority of experiments. C.S.P analyzed and interpreted metabolic data. M.D ran the lectin array experiments. E.P, R.J, K.M, J.R.K, P.T, A.L, L.J.M, J.J, and J.Z.L selected study participants and interpreted clinical data. A.R.G and H.T performed metabolic analysis. X.Y and Q.L performed statistical analysis for the whole study. L.B.G, C.S.P, and M.A-M wrote the manuscript, and all authors edited it.

## ACKNOWLEDGMENTS

This work is supported by the Foundation for AIDS Research (amfAR) impact grant # 109840-65-RGRL to M.A-M and J.Z.L as well as the NIH R21 AI143385 to M.A-M. M.A-M is also supported by NIH grants (R01 DK123733, R01 AG062383, R01NS117458, R21 AI129636, and R21 NS106970), the Penn Center for AIDS Research (P30 AI 045008), and W.W. Smith Charitable Trust grant # A1901. L.J.M is supported by R01AI48398, the NIH-funded BEAT-HIV Martin Delaney Collaboratory to cure HIV-1 infection (1UM1Al126620), Kean Family Professorship, and the Roberts I. Jacobs Fund of the Philadelphia Foundation. Metabolomics analysis was performed by the Wistar Proteomics and Metabolomics Shared Resource supported in part by NIH Cancer Center Support Grant CA010815 on a Thermo Q-Exactive HF-X mass spectrometer purchased with NIH grant S10 OD023586. This work was also supported by the National Institutes of Health (NIH) grant UM1 AI068634 to the Statistical and Data Management Center of the AIDS Clinical Trials Group, UM1 AI068636 to AIDS Clinical Trials Group, and a subcontract from UM1 AI106701 to the Harvard Virology Support Laboratory. We would like to thank Rachel E. Locke, Ph.D., for providing comments. We would like to thank all donor participants. We thank the participants, staff, and principal investigators of the ACTG studies A371, A5024, A5068, A5170, A5187 and A5197.

