## Supplementary figures and images for "Non-Invasive Plasma Glycomic and Metabolic Biomarkers of Post-treatment Control of HIV"

### Supplementary Figure 1

Supplementary Figure 1

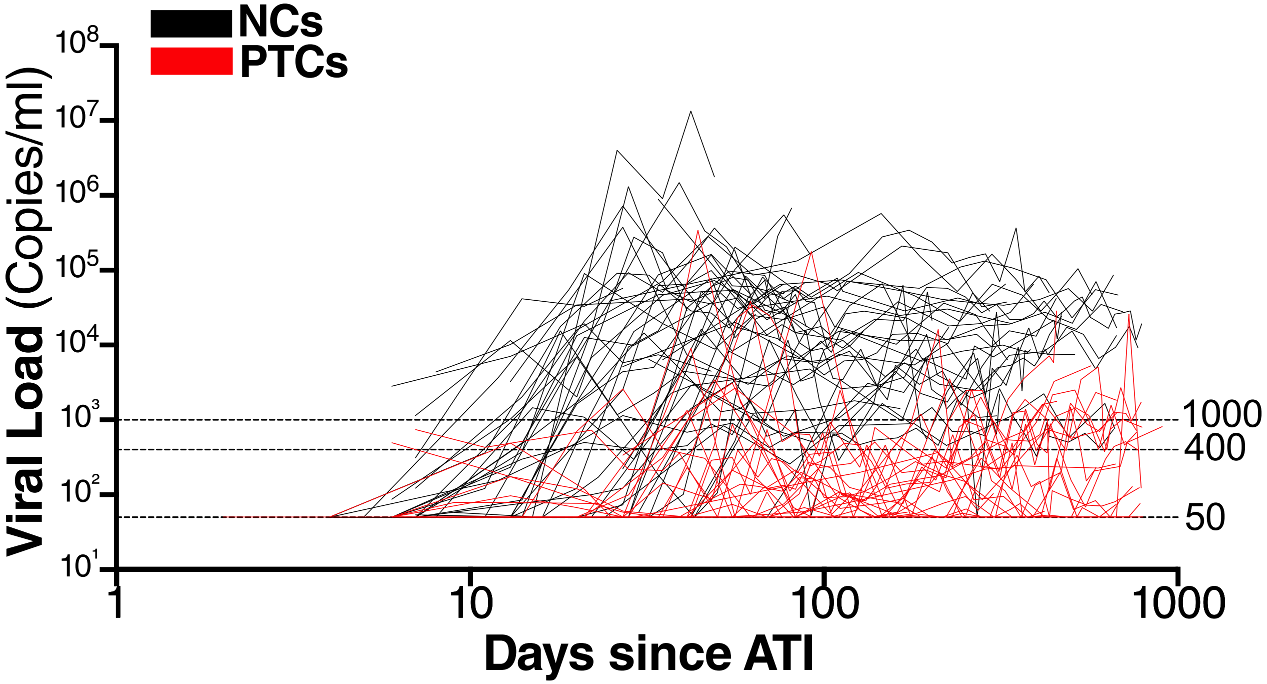

### Supplementary Figure 2

Supplementary Figure 2

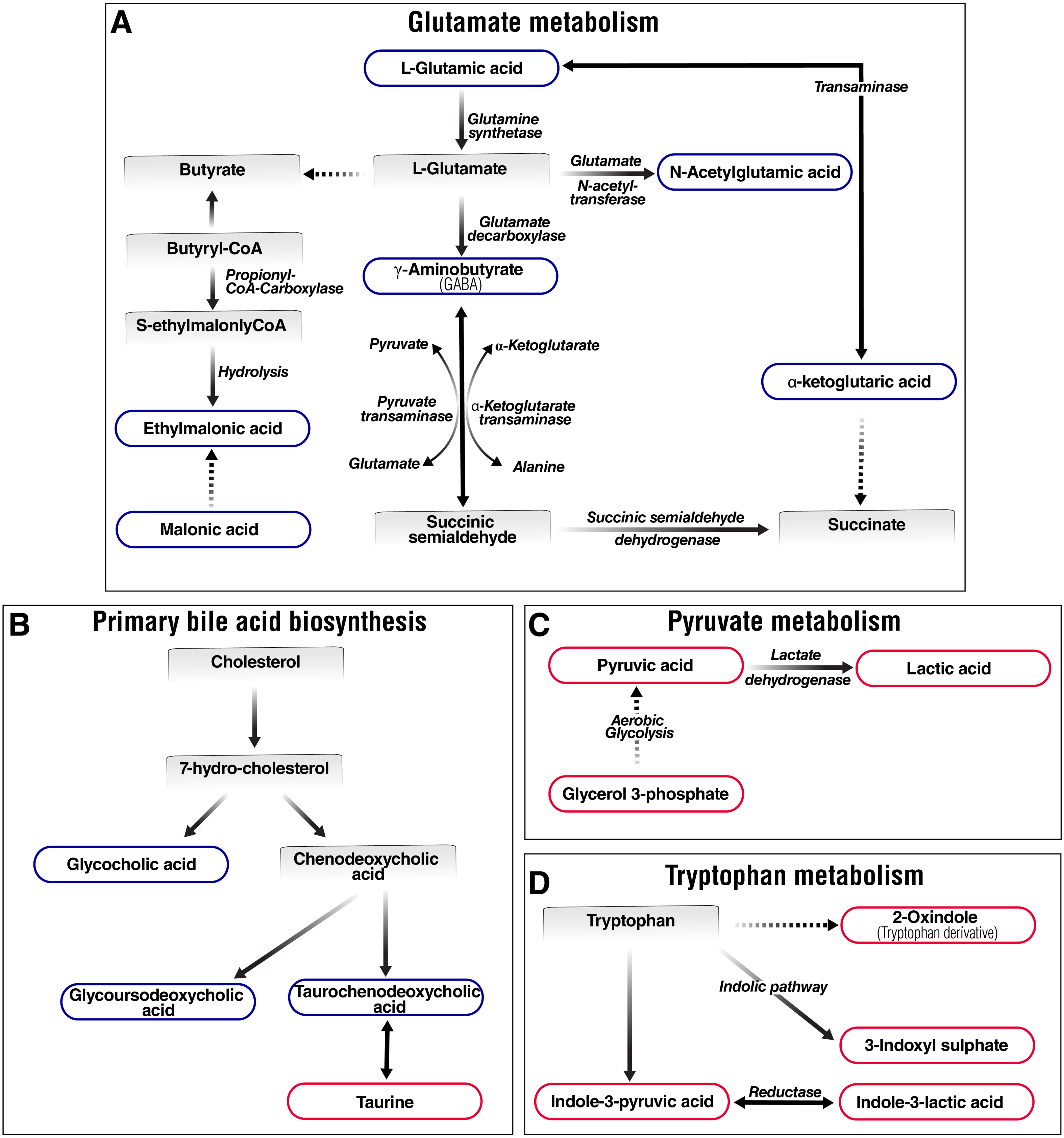

### Supplementary Figure 3

Supplementary Figure 3

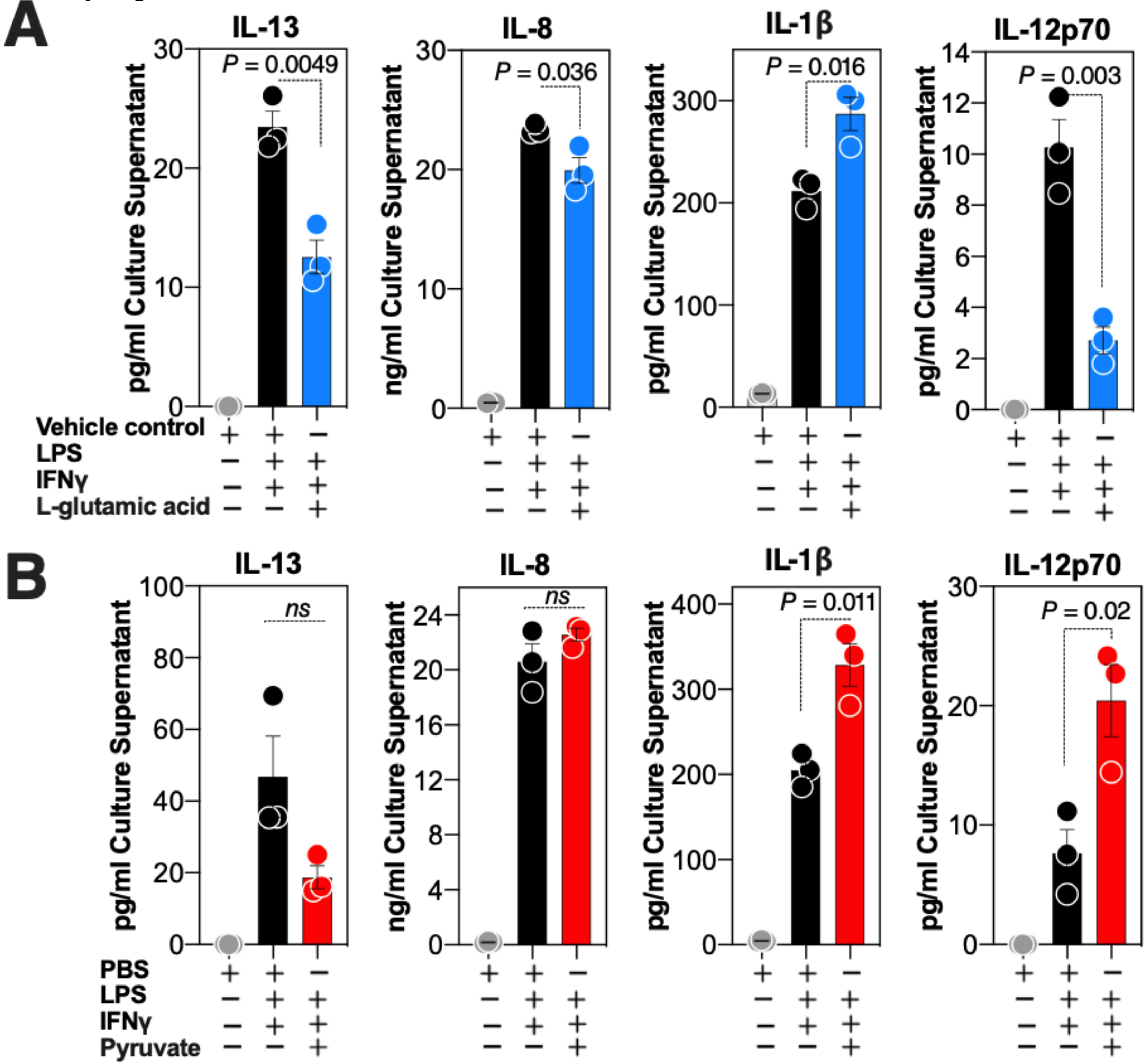

### Supplementary Figure 4

Supplementary Figure 4

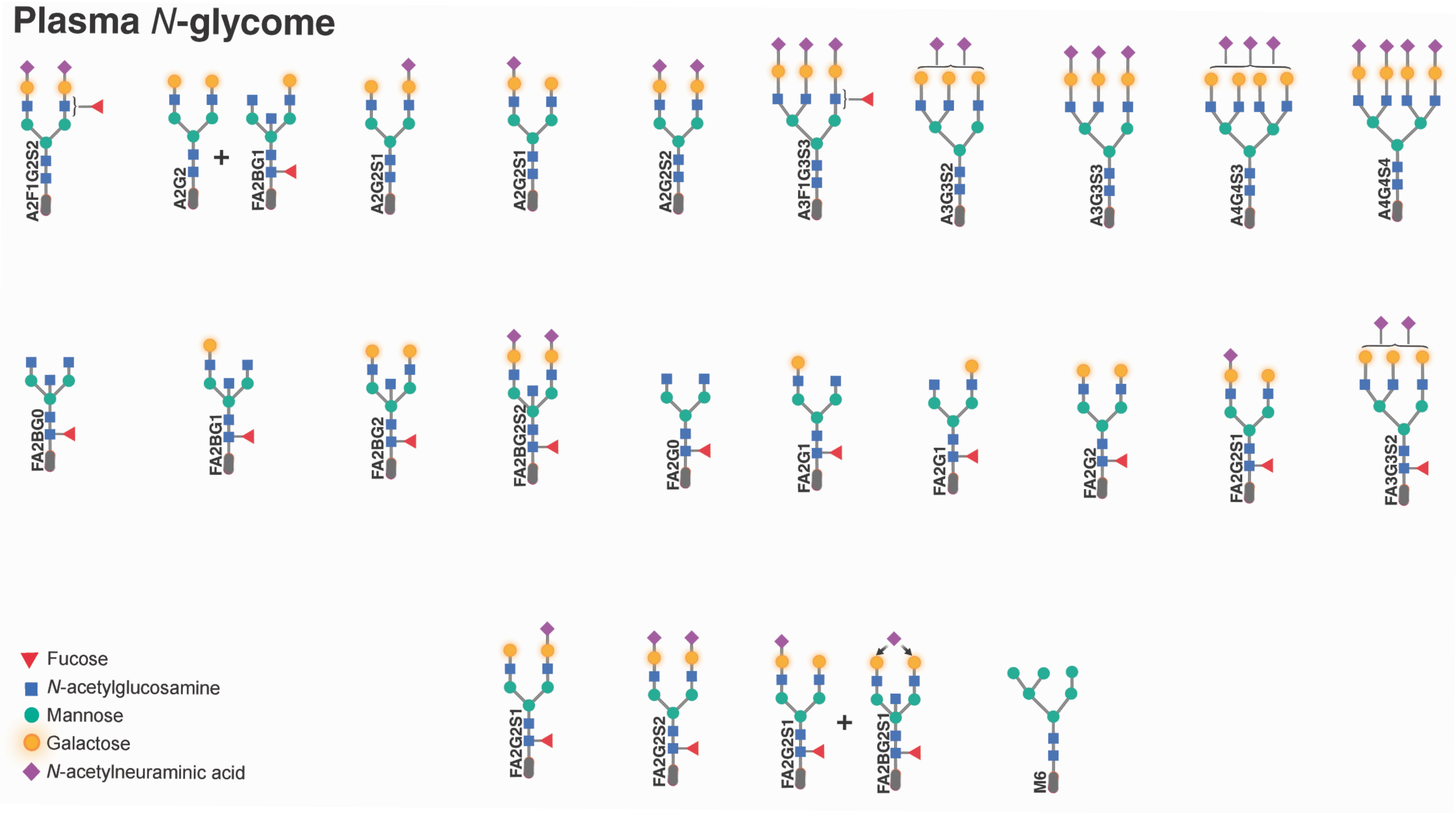
