## Supplementary Table 1 for "Non-Invasive Plasma Glycomic and Metabolic Biomarkers of Post-treatment Control of HIV"

**Supplementary Table 1.** Demographic characteristics of the Philadelphia cohort

| Patient ID | Age (years) | Gender | Race | Pre-ATI viral load (copies/ml) | Pre-ATI CD4 count (cells/mm$^3$) | Pre-ATI CD4% | Post-ATI viral setpoint (copies/ml) | Time to viral rebound (days) |
|---|---|---|---|---|---|---|---|---|
| S-8 | 53 | Male | Caucasian | <50 | 426 | 23.7 | NA | 14 |
| S-14 | 29 | Male | Caucasian | <50 | 782 | 46 | 16153 | 28 |
| S-20 | 23 | Male | African American | <50 | 708 | 37.3 | 3961 | 28 |
| S-21 | 40 | Male | Caucasian | <50 | 509 | 28.3 | 31674 | 14 |
| S-25 | 36 | Female | African American | <50 | 886 | 40.3 | 2693 | 70 |
| S-28 | 48 | Male | Hispanic | <50 | 478 | 47.8 | 274 | 28 |
| S-29 | 45 | Male | African American | <50 | 513 | 15.1 | NA | 28 |
| S-30 | 45 | Male | Caucasian | <50 | 602 | 24.1 | NA | 14 |
| S-31 | 45 | Male | Caucasian | <50 | 712 | 28.5 | 6775 | 27 |
| S-32 | 41 | Male | Caucasian | <50 | 864 | 37.6 | 21961 | 35 |
| S-33 | 49 | Female | African American | <50 | 901 | 39.2 | NA | 15 |
| S-35 | 51 | Male | Caucasian | <50 | 516 | 21.5 | 7260 | 14 |
| S-36 | 40 | Male | Caucasian | <50 | 517 | 39.8 | NA | 35 |
| S-38 | 42 | Male | Caucasian | <50 | 688 | 32.8 | 8460 | 21 |
| S-41 | 54 | Male | Caucasian | <50 | 766 | 36.5 | 21246 | 28 |
| S-42 | 42 | Male | Caucasian | <50 | 734 | 36.7 | NA | 14 |
| S-43 | 50 | Male | Caucasian | <50 | 690 | 23.8 | 13675 | 28 |
| S-44 | 44 | Male | Caucasian | <50 | 374 | 18.7 | 81949 | 14 |
| S-49 | 54 | Male | African American | <50 | 557 | 42.9 | NA | 119 |
| S-51 | 53 | Male | Caucasian | <50 | 434 | 36.2 | 1297 | 14 |
| S-54 | 43 | Male | Caucasian | <50 | 832 | 34.7 | 21456 | 14 |
| S-55 | 38 | Male | African American | <50 | 615 | 32.4 | NA | 42 |
| S-60 | 51 | Male | Caucasian | <50 | 669 | 39.4 | 16249 | 21 |
| S-62 | 48 | Male | African American | <50 | 720 | 42.4 | 6423 | 42 |
