## Supplementary Table 2 for "Non-Invasive Plasma Glycomic and Metabolic Biomarkers of Post-treatment Control of HIV"

**Supplementary Table 2.** A list of metabolites their pre-ATI levels associate with time-to-viral-rebound upon ART cessation

| Metabolites Associate with Delayed Viral Rebound | | Cox proportional-hazards model | | | | | Mantel-Cox Test | |
|---|---|---|---|---|---|---|---|---|
| **Metabolite** | **Pathway** | **HazardRatio (HR)** | **HR 95% lower confidence limit (LCL)** | **HR 95% higher confidence limit (HCL)** | ***P* value** | ***FDR*** | ***P* value** | ***FDR*** |
| **L-glutamic acid** | Glutamate metabolism | 0.157 | 0.045 | 0.555 | 0.004 | 0.072 | ns | |
| **α-ketoglutaric acid** | Glutamate metabolism | 0.181 | 0.052 | 0.626 | 0.007 | 0.089 | ns | |
| **Gamma-Aminobutyric acid** | Glutamate metabolism | 0.190 | 0.060 | 0.599 | 0.005 | 0.074 | 0.012 | 0.181 |
| **N-Acetylglutamic acid** | Glutamate metabolism | 0.355 | 0.143 | 0.881 | 0.026 | 0.158 | 0.048 | 0.296 |
| **Ethylmalonic acid** | Glutamate metabolism | 0.393 | 0.186 | 0.829 | 0.014 | 0.132 | 0.002 | 0.175 |
| **Taurochenodeoxycholic acid** | Primary bile acid biosynthesis | 0.615 | 0.450 | 0.839 | 0.002 | 0.062 | 0.007 | 0.181 |
| **Glycoursodeoxycholic acid** | Primary bile acid biosynthesis | 0.598 | 0.447 | 0.800 | 0.001 | 0.062 | 0.004 | 0.175 |
| **Glycocholic acid** | Primary bile acid biosynthesis | 0.754 | 0.625 | 0.909 | 0.003 | 0.062 | 0.025 | 0.226 |
| **D-Glucose** | Carbohydrate Metabolsim | 0.423 | 0.186 | 0.959 | 0.039 | 0.196 | 0.003 | 0.175 |
| **1,5-Anhydro-D-glucitol** | Carbohydrate Metabolsim | 0.284 | 0.087 | 0.928 | 0.037 | 0.190 | 0.009 | 0.181 |
| **Kojic acid** | Others | 0.206 | 0.073 | 0.580 | 0.003 | 0.062 | 0.016 | 0.189 |
| **Malonic acid** | Others | 0.323 | 0.117 | 0.890 | 0.029 | 0.173 | 0.039 | 0.296 |
| **D-Ribono-1,4-lactone** | Others | 0.202 | 0.054 | 0.762 | 0.018 | 0.132 | 0.011 | 0.181 |
| **Metabolites Associate with Faster Viral Rebound** | | | | | | | | |
| **Pyruvic acid** | Pyruvate metabolism | 6.793 | 2.063 | 22.362 | 0.002 | 0.062 | 0.048 | 0.296 |
| **Glycerol 3-phosphate** | Pyruvate metabolism | 3.747 | 1.561 | 8.992 | 0.003 | 0.062 | 0.048 | 0.296 |
| **L-lactic acid** | Pyruvate metabolism | 6.096 | 1.599 | 23.236 | 0.008 | 0.093 | ns | |
| **Indole-3-pyruvic acid** | Tryptophan metabolism | 2.474 | 1.170 | 5.230 | 0.018 | 0.132 | 0.031 | 0.264 |
| **Indole-3-lactic acid** | Tryptophan metabolism | 3.161 | 1.317 | 7.583 | 0.010 | 0.099 | 0.009 | 0.181 |
| **3-Indoxyl sulphate** | Tryptophan metabolism | 2.016 | 1.189 | 3.417 | 0.009 | 0.097 | 0.043 | 0.296 |
| **2-Oxindole** | Tryptophan metabolism | 2.204 | 1.358 | 3.578 | 0.001 | 0.062 | ns | |
| **Trimethylamine N-oxide** | Others/Microbiome metabolism | 1.488 | 1.123 | 1.973 | 0.006 | 0.078 | 0.008 | 0.181 |
| **Taurine** | Others/Primary bile acid biosynthesis | 4.379 | 1.561 | 12.284 | 0.005 | 0.075 | 0.002 | 0.175 |
| **Imidazolelactic acid** | Others | 2.582 | 1.403 | 4.753 | 0.002 | 0.062 | 0.020 | 0.224 |
| **Glycerophospho-N-palmitoyl ethanolamine** | Others | 2.493 | 1.265 | 4.913 | 0.008 | 0.093 | 0.044 | 0.296 |
| **Nicotinamide** | Others | 1.947 | 1.303 | 2.908 | 0.001 | 0.062 | ns | |
