## Supplementary Table 3 for "Non-Invasive Plasma Glycomic and Metabolic Biomarkers of Post-treatment Control of HIV"

**Supplementary Table 3.** Metabolic pathways associated with time-to-viral-rebound in the Philadelphia cohort

| **Pathways associated with delayed rebound** | ***P* value** |
|---|---|
| Arginine biosynthesis | 9.29E-05 |
| Butanoate metabolism | 0.000116 |
| Glutamine and glutamate metabolism | 0.000677 |
| Alanine, aspartate and glutamate metabolism | 0.000792 |
| Neomycin, kanamycin and gentamicin biosynthesis | 0.014148 |
| Arginine and proline metabolism | 0.028007 |
| Primary bile acid biosynthesis | 0.039973 |
| Nitrogen metabolism | 0.041899 |
| **Pathways associated with faster rebound** | |
| Pyruvate metabolism | 0.006521 |
| Taurine and hypotaurine metabolism | 0.04562 |
