## Supplementary Table 4 for "Non-Invasive Plasma Glycomic and Metabolic Biomarkers of Post-treatment Control of HIV"

**Supplementary Table 4.** Lectins used in the 45-plex lectin microarray

| | Name | Species | Origin | Glycan specificity[1,2] |
|---|---|---|---|---|
| 1 | LTL | *Lotus tetragonolobus* | *Natural* | Fuc ($Le^x$, $Le^y$) |
| 2 | PSA | *Pisum sativum* | *Natural* | α1-6Fuc up to biantenna |
| 3 | LCA | *Lens culinaris* | *Natural* | α1-6Fuc up to biantenna |
| 4 | UEAI | *Ulex europaeus* | Natural | α1-2Fuc |
| 5 | AOL | *Aspergillus oryzae* | Recombinant | α1-6Fuc (Core), α1-2Fuc (H), α1-3Fuc ($Le^x$), α1-3Fuc ($Le^a$) |
| 6 | AAL | *Aleuria aurantia* | Natural | α1-6Fuc (Core), α1-2Fuc (H), α1-3Fuc ($Le^x$), α1-3Fuc ($Le^a$) |
| 7 | MAL | *Maackia amurensis* | *Natural* | α2-3Sia |
| 8 | SNA | *Sambucus nigra* | Natural | α2-6Sia |
| 9 | SSA | *Sambucus sieboldiana* | Natural | α2-6Sia |
| 10 | TJAI | *Trichosanthes japonica* | Natural | α2-6Sia |
| 11 | PHAL | *Phaseolus vulgaris* | Natural | GlcNAcβ1-6Man (Tetraantenna) |
| 12 | ECA | *Erythrina cristagalli* | Natural | βGal |
| 13 | RCA120 | *Ricinus communis* | Natural | βGal |
| 14 | PHAE | *Phaseolus vulgaris* | *Natural* | bisecting GlcNAc |
| 15 | DSA | *Datura stramonium* | Natural | GlcNAcβ1-6Man (Tetraantenna) |
| 16 | GSLII | *Griffonia simplicifolia* | Natural | GlcNAcβ1-4Man |
| 17 | NPA | *Narcissus pseudonarcissus* | Natural | Manα1-3Man |
| 18 | ConA | *Canavalia ensiformis* | *Natural* | M3, Manα1-2Manα1-3(Manα1-6)Man, GlcNAcβ1-2Manα1-3(Manα1-6)Man |
| 19 | GNA | *Galanthus nivalis* | Natural | Manα1-3Man, Manα1-6Man |
| 20 | HHL | *Hippeastrum hybrid* | Natural | Manα1-3Man, Manα1-6Man |
| 21 | ACG | *Agrocybe cylindracea* | Natural | α2-3Sia |
| 22 | TxLcl | *Tulipa gesneriana* | *Natural* | Galactosylated N-glycans up to triantenna |
| 23 | BPL | *Bauhinia purpurea alba* | Natural | Galβ1-3GlcNAc(GalNAc), α/βGalNAc |
| 24 | TJAII | *Trichosanthes japonica* | Natural | α1-2Fuc |
| 25 | EEL | *Euonymus europaeus* | *Natural* | αGal (B) |
| 26 | ABA | *Agarics bisporus* | Natural | Galβ1-3GalNAc (T), GlcNAc |
| 27 | LEL | *Lycopersicon esculentum* | *Natural* | Polylactosamine, (GlcNAc)n |
| 28 | STL | *Solanum tuberosum* | Natural | Polylactosamine, (GlcNAc)n |
| 29 | UDA | *Urtica dioica* | Natural | (GlcNAc)n |
| 30 | PWM | *Phytolacca americana* | Natural | (GlcNAc)n |
| 31 | Jacalin | *Artocarpus integrifolia* | *Natural* | Galβ1-3GalNAc (T), GalNAcα (Tn) |
| 32 | PNA | *Arachis hypogaea* | Natural | Galβ1-3GalNAc (T) |
| 33 | WFA | *Wisteria floribunda* | *Natural* | Terminal GalNAc, LacDiNAc |
| 34 | ACA | *Amaranthus caudatus* | Natural | Galβ1-3GalNAc (T), GalNAca (Tn) |
| 35 | MPA | *Maclura pomifera* | Natural | Galβ1-3GalNAc (T), GalNAca (Tn) |
| 36 | HPA | *Helix pomatia* | *Natural* | αGalNAc (A, Tn) |
| 37 | VVA | *Vicia villosa* | *Natural* | α,βGalNAc (A, Tn, LacDiNAc) |
| 38 | DBA | *Dolichos biflorus* | Natural | α,βGalNAc (A, Tn, LacDiNAc) |
| 39 | SBA | *Glycine max* | Natural | α,βGalNAc (A, Tn, LacDiNAc) |
| 40 | Calsepa | *Calystegia sepium* | *Natural* | Biantenna with bisecting GlcNAc |
| 41 | PTL I | *Psophocarpus tetragonolobu* | *Natural* | αGalNAc (A, Tn) |
| 42 | MAH | *Maackia amurensis* | Natural | α2-3Sia |
| 43 | WGA | *Triticum vulgaris* | *Natural* | (GlcNAc)n, polySia |
| 44 | GSLIA4 | *Griffonia simplicifolia* | Natural | αGalNAc (A, Tn) |
| 45 | GSLIB4 | *Griffonia simplicifolia* | *Natural* | αGal (B) |

[1]Abbreviations: Gal (D-galactose), GalNAc (N-acetyl-galactosamine), GlcNAc (N-acetyl-glucosamine), Fuc (L-fucose), Glc (D-glucose), Sia (Sialic acid), LacNAc (N-acetyl-lactosamine).

[2]Specificity data was obtained by frontal affinity chromatography and glycoconjugate microarray.
