## Supplementary Table 5 for "Non-Invasive Plasma Glycomic and Metabolic Biomarkers of Post-treatment Control of HIV"

**Supplementary Table 5.** Glycomic and metabolic predictors of time-to-viral-rebound using samples from the ACTG Cohort after correction for potential confounders

| Adjusted for: | | | | Age | | Gender | | Ethnicity | | ART initiation (early vs. chronic tx) | | ART duration | | Pre-ATI CD4 count | |
|---|---|---|---|---|---|---|---|---|---|---|---|---|---|---|---|
| | **Variable** | **Category** | **Group** | **HR** | ***P* value** | **HR** | ***P* value** | **HR** | ***P* value** | **HR** | ***P* value** | **HR** | ***P* value** | **HR** | ***P* value** |
| **Time to HIV RNA value ≥1000 copies/mL** | **A3G3S3** | Plasma *N*-glycans | All | 3.01 | 0.001 | 2.59 | 0.010 | 3.25 | 0.001 | 3.02 | 0.001 | 3.10 | 0.001 | 2.97 | 0.001 |
| | **Pyruvic acid** | Metabolites | PTC | 2.60 | 0.014 | 2.60 | 0.012 | 2.65 | 0.014 | 2.82 | 0.011 | 2.63 | 0.012 | 2.70 | 0.023 |
| | **ConA-binding glycans** | Plasma total glycans | All | 2.56 | 0.009 | 2.45 | 0.008 | 2.52 | 0.008 | 2.64 | 0.007 | 2.57 | 0.008 | 2.45 | 0.011 |
| | **MPA-binding glycans** | Plasma total glycans | All | 2.27 | 0.0002 | 2.38 | 0.0001 | 2.28 | 0.0003 | 2.28 | 0.0002 | 2.28 | 0.0003 | 2.28 | 0.0003 |
| | **ACA-binding glycans** | Plasma total glycans | All | 2.27 | 0.006 | 2.09 | 0.011 | 2.34 | 0.004 | 2.24 | 0.006 | 2.22 | 0.007 | 2.20 | 0.008 |
| | **ACG-binding glycans** | Plasma total glycans | All | 0.53 | 0.002 | 0.57 | 0.006 | 0.54 | 0.001 | 0.53 | 0.002 | 0.54 | 0.002 | 0.53 | 0.001 |
| | **RCA120-binding glycans** | Plasma total glycans | All | 0.51 | 0.005 | 0.52 | 0.006 | 0.50 | 0.002 | 0.51 | 0.004 | 0.51 | 0.003 | 0.51 | 0.006 |
| | **AAL-binding glycans** | Plasma total glycans | All | 0.37 | 0.0001 | 0.33 | 0.00002 | 0.37 | 0.0002 | 0.39 | 0.0001 | 0.39 | 0.0001 | 0.39 | 0.0001 |
| | **DSA-binding glycans** | Plasma total glycans | All | 0.30 | 0.003 | 0.37 | 0.025 | 0.26 | 0.001 | 0.30 | 0.003 | 0.30 | 0.004 | 0.28 | 0.002 |
| | **L-glutamic acid** | Metabolites | All | 0.25 | 0.004 | 0.28 | 0.009 | 0.23 | 0.002 | 0.23 | 0.004 | 0.25 | 0.004 | 0.26 | 0.005 |
| | **G2** | IgG *N*-glycans | All | 0.19 | 0.038 | 0.19 | 0.021 | 0.22 | 0.049 | 0.22 | 0.047 | 0.22 | 0.05 | 0.21 | 0.040 |
| | **UDA-binding glycans** | Plasma total glycans | All | 0.18 | 0.001 | 0.20 | 0.002 | 0.18 | 0.001 | 0.16 | 0.0004 | 0.17 | 0.001 | 0.18 | 0.0004 |
| | **STL-binding glycans** | Plasma total glycans | All | 0.17 | 0.003 | 0.17 | 0.002 | 0.17 | 0.003 | 0.16 | 0.002 | 0.15 | 0.002 | 0.16 | 0.003 |
| **Time to two consecutive HIV RNA values ≥1000 copies/mL** | **A3G3S3** | Plasma *N*-glycans | All | 4.04 | 0.000 | 3.56 | 0.001 | 3.92 | 0.000 | 3.77 | 0.000 | 3.71 | 0.000 | 3.65 | 0.000 |
| | **ConA-binding glycans** | Plasma total glycans | All | 4.14 | 0.000 | 3.55 | 0.001 | 3.79 | 0.000 | 4.06 | 0.000 | 3.74 | 0.000 | 3.67 | 0.000 |
| | **Nicotinamide 1-oxide** | Metabolites | PTC | 4.64 | 0.012 | 3.43 | 0.030 | 3.26 | 0.037 | 3.53 | 0.029 | 3.52 | 0.032 | 3.35 | 0.033 |
| | **MPA-binding glycans** | Plasma total glycans | All | 2.86 | 0.0000 | 2.71 | 0.0000 | 2.81 | 0.0000 | 2.71 | 0.0000 | 2.72 | 0.0000 | 2.75 | 0.0000 |
| | **ACA-binding glycans** | Plasma total glycans | All | 2.23 | 0.016 | 1.97 | 0.036 | 2.10 | 0.023 | 2.07 | 0.025 | 2.03 | 0.028 | 2.03 | 0.031 |
| | **ABA-binding glycans** | Plasma total glycans | All | 2.06 | 0.022 | 2.19 | 0.013 | 2.08 | 0.019 | 2.06 | 0.021 | 2.04 | 0.025 | 2.16 | 0.016 |
| | **α-ketoglutaric acid** | Metabolites | All | 0.46 | 0.001 | 0.54 | 0.002 | 0.51 | 0.001 | 0.48 | 0.001 | 0.54 | 0.003 | 0.54 | 0.002 |
| | **LCA-binding glycans** | Plasma total glycans | All | 0.52 | 0.042 | 0.48 | 0.028 | 0.53 | 0.048 | 0.49 | 0.031 | 0.52 | 0.037 | 0.48 | 0.023 |
| | **ACG-binding glycans** | Plasma total glycans | All | 0.48 | 0.000 | 0.53 | 0.001 | 0.52 | 0.001 | 0.51 | 0.001 | 0.52 | 0.001 | 0.51 | 0.000 |
| | **DSA-binding glycans** | Plasma total glycans | All | 0.30 | 0.004 | 0.35 | 0.021 | 0.30 | 0.005 | 0.32 | 0.007 | 0.32 | 0.009 | 0.28 | 0.003 |
| | **AAL-binding glycans** | Plasma total glycans | All | 0.28 | 0.000 | 0.25 | 0.000 | 0.25 | 0.000 | 0.28 | 0.000 | 0.29 | 0.000 | 0.28 | 0.000 |
| | **L-glutamic acid** | Metabolites | All | 0.23 | 0.006 | 0.25 | 0.010 | 0.22 | 0.004 | 0.22 | 0.006 | 0.24 | 0.01 | 0.24 | 0.009 |
| | **G2** | IgG *N*-glycans | All | 0.11 | 0.005 | 0.14 | 0.009 | 0.15 | 0.013 | 0.15 | 0.0144 | 0.15 | 0.019 | 0.14 | 0.0119 |
| | **UDA-binding glycans** | Plasma total glycans | All | 0.12 | 0.000 | 0.14 | 0.000 | 0.13 | 0.000 | 0.11 | 0.000 | 0.13 | 0.000 | 0.13 | 0.000 |
| | **STL-binding glycans** | Plasma total glycans | All | 0.12 | 0.001 | 0.12 | 0.001 | 0.11 | 0.001 | 0.12 | 0.001 | 0.11 | 0.001 | 0.11 | 0.001 |

HR = Hazard Ratio
