## Supplementary Table 6 for "Non-Invasive Plasma Glycomic and Metabolic Biomarkers of Post-treatment Control of HIV"

**Supplementary Table 6.** Comparisons of C-index between each univariate Cox model versus multivariable Cox model with Lasso selected variables predict time to VL ≥1000

| Predictors in the model | N | C-index | SE | 95% Confidence interval | | *P*-value (single predictor models vs. Lasso selected multivariable model) |
|---|---|---|---|---|---|---|
| Variables identified by Lasso multivariable Cox model | 70 | 0.74 | 0 | 0.7 | 0.8 | reference |
| **Plasma A3G3S3 glycans** | 70 | 0.64 | 0 | 0.6 | 0.71 | 0.017 |
| **Pyruvic acid** | 70 | 0.56 | 0 | 0.5 | 0.64 | <0.0001 |
| Plasma mannose (ConA binding) | 70 | 0.58 | 0 | 0.5 | 0.66 | 0.001 |
| **Plasma T-antigen (MPA binding)** | 70 | 0.62 | 0 | 0.5 | 0.7 | 0.007 |
| **Plasma T-antigen (ACA binding)** | 70 | 0.62 | 0 | 0.5 | 0.69 | <0.0001 |
| α2,3 sialic acid (ACG binding) | 70 | 0.49 | 0.1 | 0.4 | 0.58 | <0.0001 |
| βGal (RCA120 binding) | 70 | 0.55 | 0.1 | 0.5 | 0.64 | <0.0001 |
| **Total fucose (AAL binding)** | 70 | 0.62 | 0 | 0.5 | 0.71 | 0.004 |
| GlcNAcβ1-6Mannose (DSA binding) | 70 | 0.53 | 0 | 0.5 | 0.61 | <0.0001 |
| **L-glutamic acid** | 70 | 0.63 | 0 | 0.6 | 0.7 | 0.001 |
| IgG G2 glycans | 70 | 0.54 | 0.1 | 0.4 | 0.63 | <0.0001 |
| (GlcNAc)n (UDA binding) | 70 | 0.6 | 0 | 0.5 | 0.68 | 0.004 |
| **(GlcNAc)n (STL binding)** | 70 | 0.63 | 0 | 0.6 | 0.7 | 0.003 |

Bold variables are those selected by Lasso to be included in the multivariable Cox model predicting time to VL ≥1000. C-index = The concordance index; SE = Standard error
