## Supplementary Table 7 for "Non-Invasive Plasma Glycomic and Metabolic Biomarkers of Post-treatment Control of HIV"

**Supplementary Table 7.** Comparisons of C-index between each univariate Cox model versus multivariable Cox model with Lasso selected variables predict time to two consecutive VL ≥1000

| **Predictors in the model** | **N** | **C-index** | **SE** | **95% Confidence interval** | | ***P*-value (single predictor models vs. Lasso selected multivariable model** |
|---|---|---|---|---|---|---|
| Variables identified by Lasso multivariable Cox model | 70 | 0.764 | 0.039 | 0.7 | 0.842 | reference |
| **Plasma A3G3S3 glycans** | 70 | 0.651 | 0.039 | 0.6 | 0.729 | 0.022 |
| **Plasma mannose (ConA binding)** | 70 | 0.599 | 0.045 | 0.5 | 0.688 | 0.004 |
| **Nicotinamide1 oxide** | 70 | 0.532 | 0.04 | 0.5 | 0.611 | <0.0001 |
| Plasma T-antigen (MPA binding) | 70 | 0.623 | 0.044 | 0.5 | 0.711 | 0.002 |
| **Plasma T-antigen (ACA binding)** | 70 | 0.605 | 0.04 | 0.5 | 0.685 | 0.001 |
| **Plasma T-antigen (ABA binding)** | 70 | 0.631 | 0.043 | 0.5 | 0.718 | 0.002 |
| **α-ketoglutaric acid** | 70 | 0.623 | 0.041 | 0.5 | 0.706 | 0.006 |
| Core fucose (LCA binding) | 70 | 0.574 | 0.045 | 0.5 | 0.663 | <0.0001 |
| **α2,3 sialic acid (ACG binding)** | 70 | 0.508 | 0.046 | 0.4 | 0.601 | <0.0001 |
| **GlcNAcβ1-6Mannose (DSA binding)** | 70 | 0.51 | 0.041 | 0.4 | 0.591 | <0.0001 |
| **Total fucose (AAL binding)** | 70 | 0.649 | 0.042 | 0.6 | 0.733 | 0.003 |
| **L-glutamic acid** | 70 | 0.633 | 0.038 | 0.6 | 0.708 | 0.002 |
| **IgG G2 glycans** | 70 | 0.546 | 0.051 | 0.4 | 0.648 | <0.0001 |
| (GlcNAc)n (UDA binding) | 70 | 0.605 | 0.044 | 0.5 | 0.692 | 0.001 |
| **(GlcNAc)n (STL binding)** | 70 | 0.641 | 0.041 | 0.6 | 0.723 | 0.006 |

Bold variables are those selected by Lasso to be included in the multivariable Cox model predicting time to two consecutive VL ≥1000. C-index = The concordance index; SE = Standard error
