## Supplementary Table 8 for "Non-Invasive Plasma Glycomic and Metabolic Biomarkers of Post-treatment Control of HIV"

**Supplementary Table 8.** Glycomic and metabolic Variables differentiating PTCs from NCs using samples from the ACTG Cohort after correction for potential confounders

| | Adjusted for: | | Age | Gender | Ethnicity | ART initiation (early vs. chronic tx) | ART duration | Pre-ATI CD4 count |
|---|---|---|---|---|---|---|---|---|
| **Variable** | **Category** | **Comparisons** | ***P* value** | ***P* value** | ***P* value** | ***P* value** | ***P* value** | ***P* value** |
| **A2** | IgG *N*-glycans | PTC vs NC | 0.025 | 0.019 | 0.021 | 0.008 | 0.019 | 0.018 |
| **A3G3S3** | Plasma *N*-glycans | PTC vs NC | 0.0001 | 0.0004 | 0.0003 | 0.011 | 0.0002 | 0.0003 |
| **AAL-binding glycans** | Plasma total glycans | PTC vs NC | 0.00001 | 0.00001 | 0.00002 | 0.00001 | 0.00001 | 0.00001 |
| **PSA-binding glycans** | Plasma total glycans | PTC vs NC | 0.004 | 0.004 | 0.004 | 0.003 | 0.003 | 0.002 |
| **LCA-binding glycans** | Plasma total glycans | PTC vs NC | 0.004 | 0.003 | 0.003 | 0.002 | 0.003 | 0.001 |
| **ABA-binding glycans** | Plasma total glycans | PTC vs NC | 0.008 | 0.003 | 0.003 | 0.003 | 0.004 | 0.003 |
| **STL-binding glycans** | Plasma total glycans | PTC vs NC | 0.0002 | 0.001 | 0.001 | 0.001 | 0.001 | 0.000 |
| **UDA-binding glycans** | Plasma total glycans | PTC vs NC | 0.001 | 0.003 | 0.002 | 0.002 | 0.002 | 0.001 |
| **α-ketoglutaric acid** | Metabolites | PTC vs NC | 0.002 | 0.010 | 0.007 | 0.019 | 0.010 | 0.010 |
| **L-glutamic acid** | Metabolites | PTC vs NC | 0.008 | 0.011 | 0.006 | 0.000 | 0.007 | 0.010 |
