## Supplementary Table 9 for "Non-Invasive Plasma Glycomic and Metabolic Biomarkers of Post-treatment Control of HIV"

**Supplementary Table 9.** Comparisons of AUC between each logistic model with single predictor versus multivariable logistic regression with lasso selected variables predicting outcome (NC vs. PTC)

| Predictors in the model | N | AUC | SE | 95% Confidence interval | | P-value (single predictor models vs. Lasso selected multivariable model) |
|---|---|---|---|---|---|---|
| Variables identified by Lasso multivariable logistic model | 70 | 0.98 | 0.016 | 0.94 | 1 | reference |
| **IgG A2 glycans** | 70 | 0.87 | 0.039 | 0.8 | 0.95 | 0.0013 |
| **Plasma A3G3S3 glycans** | 70 | 0.72 | 0.068 | 0.59 | 0.85 | 0.0001 |
| **Total fucose (AAL binding)** | 70 | 0.79 | 0.053 | 0.69 | 0.89 | 0.0003 |
| Core fucose (PSA binding) | 70 | 0.69 | 0.064 | 0.57 | 0.82 | <0.0001 |
| **Core fucose (LCA binding)** | 70 | 0.71 | 0.063 | 0.58 | 0.83 | <0.0001 |
| **Plasma T-antigen (ABA binding)** | 70 | 0.75 | 0.063 | 0.63 | 0.87 | 0.0001 |
| **(GlcNAc)n (STL binding)** | 70 | 0.8 | 0.06 | 0.68 | 0.91 | 0.0018 |
| (GlcNAc)n (UDA binding) | 70 | 0.72 | 0.066 | 0.6 | 0.85 | <0.0001 |
| **L-glutamic acid** | 70 | 0.69 | 0.064 | 0.57 | 0.82 | <0.0001 |
| α-ketoglutaric acid | 70 | 0.67 | 0.071 | 0.53 | 0.81 | <0.0001 |

Bold variables are those selected by Lasso to be included in the multivariable logistic model predicting PTC status (PVR score); AUC = Area under the ROC Curve; SE = Standard error
